# The invasion phenotypes of glioblastoma depend on plastic and reprogrammable cell states

**DOI:** 10.1101/2024.04.23.589925

**Authors:** Milena Doroszko, Rebecka Stockgard, Irem Yildirim, Mar Ballester Bravo, Thomas O. Millner, Soumi Kundu, Josephine Heinold, Ramy Elgendy, Maria Berglund, Ludmila Elfineh, Cecilia Krona, Silvia Marino, Ida Larsson, Sven Nelander

**Affiliations:** Department of Immunology, Genetics and Pathology, Uppsala University, Program for Neurooncology and Neurodegeneration, Uppsala, SE-752 81, Sweden; Department of Pediatric Oncology, Dana-Farber Cancer Institute, Boston, MA, 02215-5450, United States; Brain Tumour Research Centre, Blizard Institute, Faculty of Medicine and Dentistry, Queen Mary University of London, London E1 4NS, United Kingdom

## Abstract

Glioblastoma (GBM), the most common primary brain cancer in adults, is characterized by rapid local invasion along diverse routes, such as infiltration of white matter tracts and penetration of perivascular spaces. We investigate the hypothesis that GBM invasion routes correlate with the transcriptional states of individual cells and identify regulators of route-specific invasion. Utilizing patient-derived GBM xenograft models, we integrate single-cell transcriptomics and spatial proteomics, revealing that mesenchymal and oligodendrocyte progenitor-like GBM cells migrate perivascularly, while neural progenitor and astrocyte-like GBM cells invade diffusely. Computational reconstruction identifies *ANXA1* as a perivascular invasion driver and lineage-restricted transcription factors *RFX4* and *HOPX* as drivers of diffuse invasion, predictive of patient survival. Genetic ablation of these genes alters invasion phenotypes and extends survival in xenografted mice, clarifying the role of cell states in GBM invasion, and highlighting potential therapeutic targets for selective invasion route targeting in GBM patients.

## Introduction

Glioblastoma (GBM), the most common primary brain cancer in adults, is characterized by rapid progression and a lack of effective therapeutic options for patients with recurrent disease. Unlike other difficult forms of cancer, GBM causes death not by distant metastasis but by rapid local invasion. The recurrence of GBM is attributed to infiltrative cells found in perivascular spaces, white matter, or brain parenchyma, also known as Secondary Scherer structures [1, 2]. The amount of infiltration is negatively correlated with overall survival and tumor growth rate, as supported by surgical [3], radiological [4], mathematical [5], and animal model studies. Yet, infiltrating cells are largely out of reach for current therapy. Comparisons between present-day patients and historical cases suggest that while the severe mass effect appears to be less common in GBM patients today, dissemination, including life-threatening brainstem invasion, is now more pronounced [6].

These observations prompt several questions. First, is the consistently observed effect of invasion on survival driven by particular types of invading cells? Second, what are the cell-intrinsic and extrinsic factors that mediate different types of invasion, and do they differ among patients? Lastly, can we kill invading cells or at least mitigate invasion to substantially prolong survival in recurrent GBM? Recent molecular studies, including single-cell profiling, indicate that GBM cells exist in transcriptionally distinct subpopulations, known as cell states, influenced by both cell-intrinsic factors (likely mutations and epigenetic factors) and the microenvironment [7–15]. According to one often used classification [8], GBM cells exist in four states termed mesenchymal-like (MES-like), oligodendrocyte precursor cell (OPC)-like, neural progenitor cell (NPC)-like, and astrocyte (AC)-like cells. Of these, the mesenchymal state has been suggested to be important for invasion [16]. Recently, the GLASS consortium reported a gradual increase of the mesenchymal subtype in patients over time, which seems consistent with an increased invasive component in recurrent tumors [17]. Intriguingly, Venkataramani et al. [18] described a model of GBM in which primarily the OPC/NPC-like states were diffusely invading *in vivo*. A growing number of pathways have been linked to invasion, such as signaling via the Ephand epidermal growth factor receptors [14, 19, 20], stemness pathways [21], and transcription factors such as *SOX10* and *CEBPB* [16, 22]. Altogether, the genetic regulation and therapeutic importance of GBM invasion processes largely remain an open problem.

Here, we test the hypothesis that the ability of cells to invade along specific routes is tightly linked to transcriptional states. In particular, we aim to understand which GBM cell states prefer invasion along the perivascular vs. diffuse route, highlight key functional properties of these states, and identify genes essential for specific types of invasion. Using patient-derived xenograft (PDX) models with a diverse spectrum of invasion patterns, we integrate single-cell transcriptomics and spatial proteomics, revealing distinctive migration patterns of GBM cell subpopulations. Our study reveals three novel genetic regulators of route-specific invasion: *ANXA1*, implicated in tumor invasion; *HOPX*, which regulates outer radial glia cells; and *RFX4*, a gene crucial for neural lineage commitment. These findings were further validated in two patient cohorts, offering promising avenues for therapeutic exploration. Jointly, our results define a tight association between cell states and GBM invasion, identify new biomarkers of route-specific invasion, and point toward targeted modulation of the cell state as a therapeutic strategy in GBM.

## Results

### HGCC xenografts display a wide range of growth structures and invasion routes

The Human Glioblastoma Cell Culture (HGCC) Resource consists of extensively studied patient-derived cell (PDC) cultures, thoroughly investigated at genomic and pharmacological levels [23, 24]. In our ongoing research, we have been systematically characterizing the invasion phenotypes of 64 GFP/luciferase-tagged HGCC cultures in nude mice, assessing the extent of perivascular and diffuse invasion, along with other morphological characteristics. The two predominant phenotypes identified in these studies are either a consolidated tumor with perivascular invasion or a diffuse growth pattern, frequently involving the corpus callosum (Krona et al., unpublished data). For this study, we selected six HGCC cell cultures that consistently displayed one of these two phenotypes. We evaluated the growth structures using multiplexed immunofluorescence staining, employing STEM121 to identify tumor cells, and specific markers such as CD31 for blood vessels, MBP for white matter, AQP4 for astrocytes, and NeuN for neurons. Three of the chosen cultures (U3013MG, U3054MG, and U3220MG) produced bulky tumors with dense perivascular growth (Figure 1A), whereas the other three (U3031MG, U3179MG, and U3180MG) produced a diffuse infiltration phenotype (Figure 1B). Several different secondary Scherer structures were evident in our models (Figure 1C), including leptomeningeal spread (U3220MG) and perineuronal satellitosis (U3031MG and U3179MG). Of note, the phenotypes demonstrated high reproducibility among mouse individuals (Supplementary figure 1, 2), with concordance levels of 96% for diffuse infiltration, 88% for perivascular invasion, and 96% for perineuronal invasion (Supplementary table 1). Interestingly, mouse survival rates varied between cases, with diffusely invading HGCC cultures showing a tendency toward longer survival times compared to those with bulk and perivascular growth and invasion phenotypes (Figure 1D, p=0.0044, log-rank test, n=45 mice). The selected cultures had a spectrum of characteristic GBM mutations (Figure 1E). In conclusion, these selected xenografts serve as representative examples of GBM with specific invasion routes.

**Figure 1:**
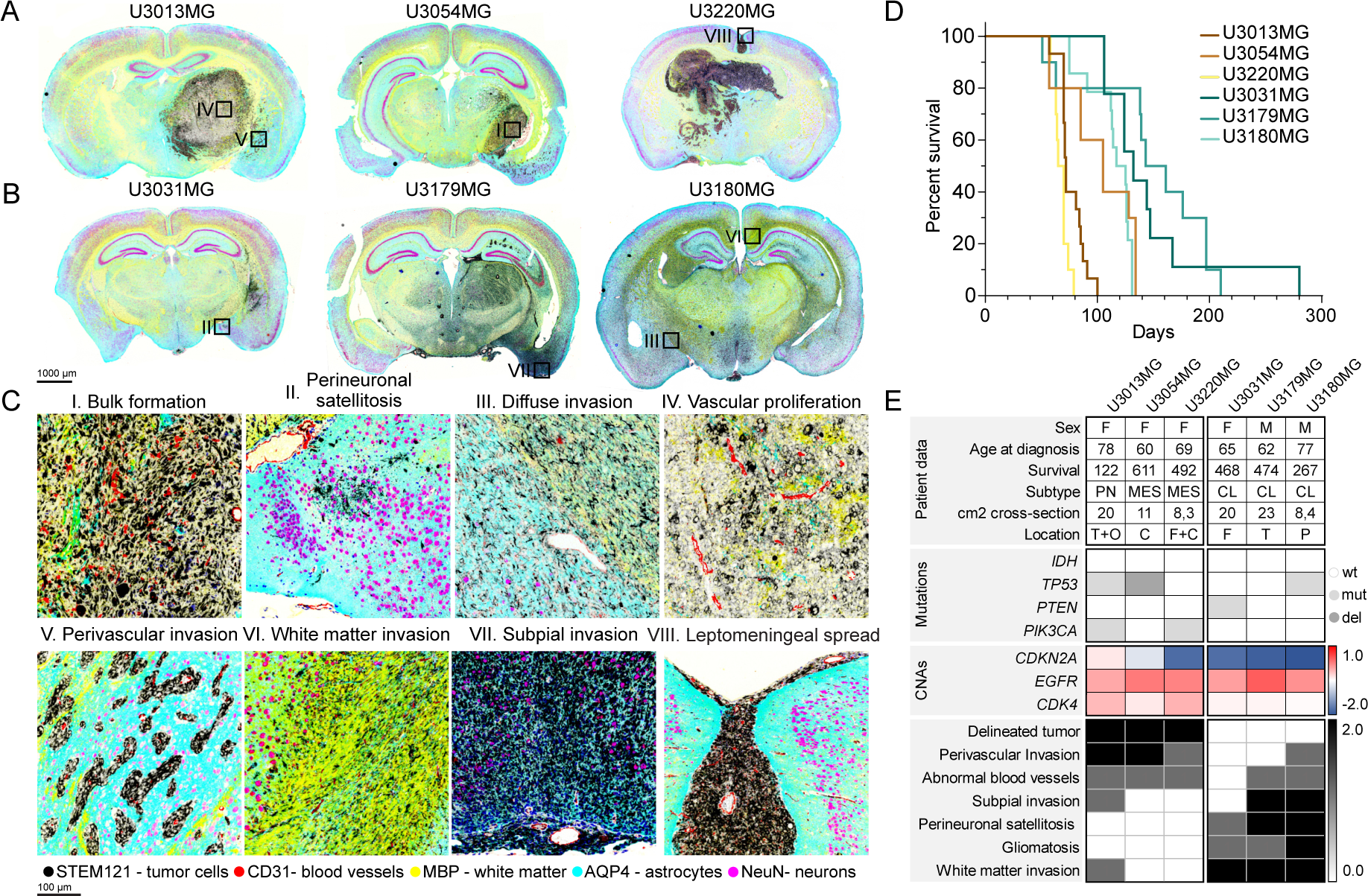
Glioblastoma xenografts recapitulate known growth patterns. (A) Coronal sections of mouse brains xenotransplanted with (A) U3013MG, U3045MG, and U3220MG PDCs (B) U3031MG, U3179MG, and U3180MG PDCs. Tumor cells are visualized in black (STEM121), blood vessels in red (CD31), white matter in yellow (MPB), astrocytes in cyan (AQP4), and neurons in magenta (NeuN). The scale bar indicates 1000 μm. Black squares indicate the zoomed-in location in (C). (C) I) Bulk formation, II) perineuronal satellitosis, III) diffuse invasion, IV) vascular proliferation, V) Perivascular invasion route, VI) White matter invasion route, VII) Subpial invasion route, VIII) Leptomeningeal spread. The scale bar indicates 100 μm. (D) Mouse survival in days for each injected PDC. (E) Table detailing patient data, mutational profiles, CNAs, observed phenotype, and invasion route.

### Transcriptional states define invasion routes in GBM xenograft models

Next, we aimed to elucidate the connection between the cell state distribution and the invasion phenotype in our PDX models, utilizing single-cell RNA sequencing (scRNA-seq) for each culture. This encompassed samples from adherent cultures before injection and tumor cells isolated from mouse brains at experimental endpoint, totaling 19 scRNA-seq samples with 119,766 cell transcriptomes passing quality control. The UMAP dimensionality reduction (Figure 2A, B) and gene set enrichment of markers obtained by graph-based cell clustering (Figure 2C) revealed distinct regions within the gene expression space for cells derived from the two classes of PDXs. Specifically, PDX models with bulk-forming and perivascular invading tumors populated a transcriptional subspace enriched for injury response and macrophage-like expression signatures, while diffusely growing PDX models occupied a region enriched for neurodevelopmental, neuronal-like signatures. Oligodendrocyte-like signatures were observed for both invasion routes. These latter were also enriched for outer radial glial cell markers and astrocytic markers (Figure 2C) [25]. PDX and PDC cells grouped together with cell cycle-related programs in a UMAP dimensionality reduction, confirming that all PDX and PDC include cycling and non-cycling cells (Figures 2A, C). In addition U3220MG displayed a specific cluster apparently associated with its high degree of leptomeningeal invasion (Figure 2A).

**Figure 2:**
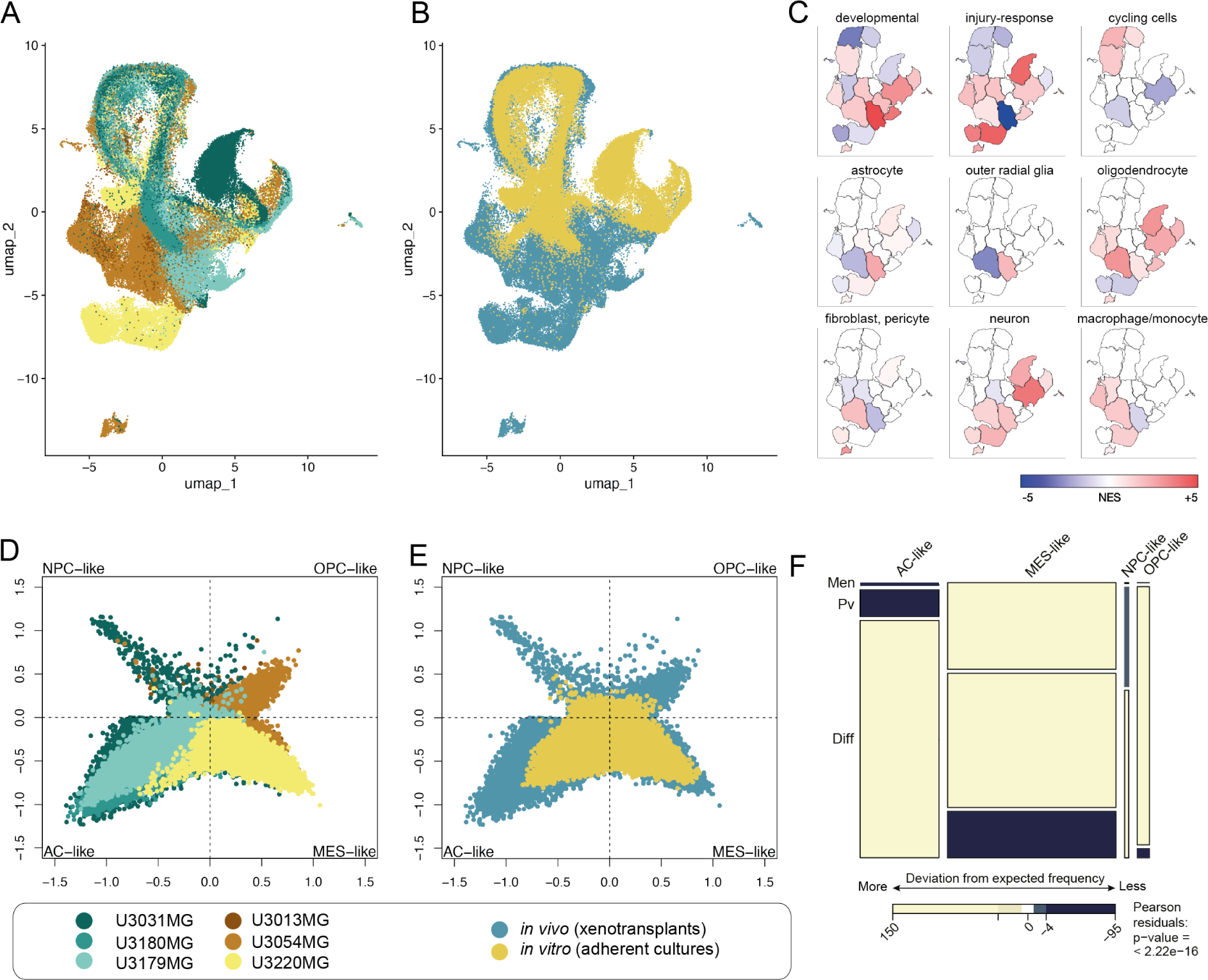
GBM cells with distinct invasion phenotypes occupy distinct transcriptional states. (A) UMAP separation of GBM cells by source patient suggests separation by invasion phenotype. (B) UMAP of the same GBM cells by growth condition, note that PDX-derived cells occupy a greater set of transcriptional states. (C) Perivascular invading cells (blue) exhibit enrichment for injury response, oligodendrocyte, and macrophage signatures, while diffusely growing cells show enrichment of neurodevelopmental signatures. (D,E) 4-state embedding (c.f. Neftel et at) shows MES/OPC enrichment of perivascular invading cells and NPC/AC enrichment of diffusely growing GBM cells. (F) Mosaic plot quantifying the relationship between transcriptional cell state and preferred invasion route, with coloring indicating observed frequencies compared to expected.

Interestingly, the cells transplanted into mice showed a wider variety of cell states compared to those cultured *in vitro* (Figures 2B, E, Supplementary figure 3). A possible explanation for this is that exposure to the mouse brain environment activates latent differentiation potential of the cells, whereas the cells stay less differentiated *in vitro*, which is maintained in stem cell conditions. We further computed cell state plots (cf. [8]), showing that the perivascular invading cultures showed a strong bias towards OPC-like and MES-like states, whereas the diffusely invading cultures were associated with NPC-like and AC-like states (pval « 0.001, Figure 2D, F). This is intriguing since a previous characterization of invasive GBM, which focused on electrophysiological connectivity of the cells, found an important separation between unconnected NPC-like and OPC-like cells on the one hand, and connected AC-like and MES-like cells on the other hand [18]. This finding suggests that the preference for perivascular vs. diffuse invasion routes is orthogonal to the electrophysiological phenotype concerning cell state.

Taken together, we found a clear correlation between the invasion patterns of PDX models and the unique cellular states they exhibited. Notably, perivascular invasion was marked by an abundance of OPC-like and MES-like states, while diffuse invasion was characterized by an NPC-like and AC-like state dominance. Of note, while this key difference was more evident in cells sampled from mouse brains, it was also seen before injection, underscoring that the tendency towards a particular invasion phenotype and cell state distribution are intrinsic properties of GBM PDCs.

### Data-driven modeling reveals potential regulators of GBM invasion routes

Our initial scRNA-seq analysis revealed a significant correlation between transcriptional states and *in vivo* invasion routes. Subsequently, we employed a data-driven approach to identify potential regulators of GBM invasion.

We have previously described a novel method, termed single-cell regulatory-driven clustering (scregclust), to simultaneously cluster genes into modules and predict regulators (such as transcription factors and kinases) of these gene modules [26] (Larsson et al., in revision, Nature Communications). Applying scregclust to the scRNA-seq data from our PDX and PDC models resulted in a regulatory landscape, where the different gene modules cluster based on their association with predicted upstream regulators (Figure 3A, Supplementary file 2). We assessed the modules by quantifying their similarity with established gene signatures of transcriptional states from [8], as well as signatures of diffuse, perivascular, and leptomeningeal invasion routes fitted from our data (Figure 3B). Upon inspection of the landscape, it became evident that groups of modules - referred to as metamodules - emerged across different PDX models, displaying shared functional profiles and regulation (Supplementary figure 4A). By projecting the metamodule gene signatures onto a single cell atlas of human cortical development [27], we could classify them according to their resemblance to normal cell types in the human brain, e.g. oligodendrocytes or astrocytes (Supplementary figure 4B). As a positive control of our regulatory predictions, we confirmed modules corresponding to the cell cycle programs G1/S and G2/M, predicted to be regulated by known cell cycle markers such as *E2F1* and *TK1* (G1/S), and *AURKA/B* (G2/M).

**Figure 3:**
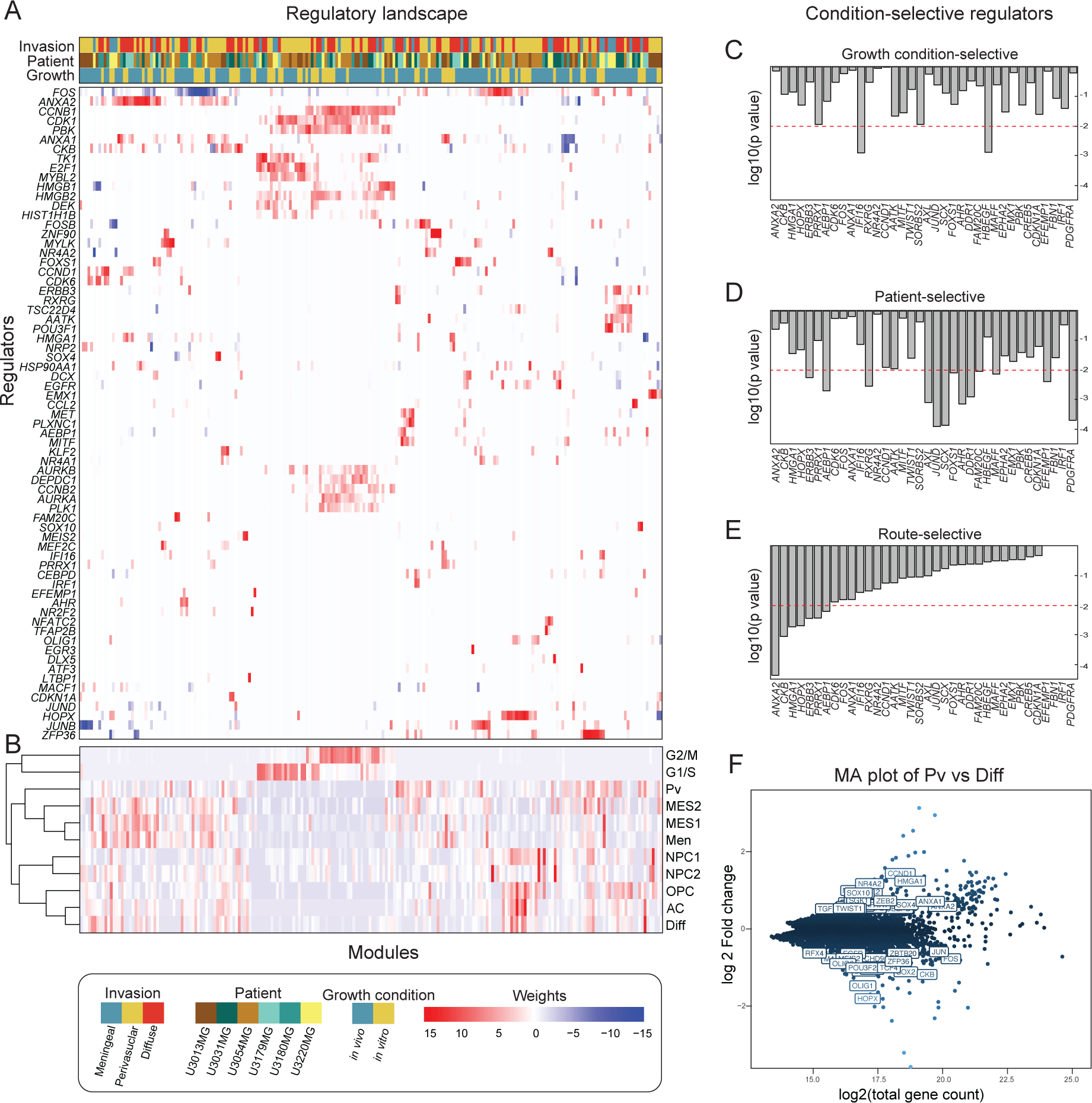
The regulatory landscape of glioblastoma cells reveals potential candidates that regulate invasion routes. (A) Heatmap of the regulatory landscape, with rows representing regulators (transcription factors and kinases) and columns representing gene modules, indicating sample origin. (B) Overlap between module gene content and cell state or invasion route signatures. (C-E) Barplots displaying regulators selective for growth condition, patient, and invasion route. (F) MA plot of differentially expressed genes, with labeled regulators from (A) with an absolute log 2 fold greater than 0.5.

We used one-way ANOVA tests to analyze how the regulators of gene modules were influenced by factors such as growth conditions (*in vitro* vs. *in vivo*) (Figure 3C), patient source (Figure 3D), or invasion routes (Figure 3E). Subsequently, we conducted a differential gene expression analysis between the perivascular and diffuse invasion routes to identify genes strongly associated with each route (Figure 3F).

Our analysis identified a total of 53 regulators associated with invasion routes: 36 linked to growth condition or source patient, and an additional 17 with differential expression between perivascular and diffuse invasion (padj<0.01, and differential expression log 2-fold change > 0.5). Key regulators attributed to the perivascular route were *ANXA1* and *ANXA2*, two members of the annexin family, that play roles in inflammation and apoptosis [28]. Regulators for the diffuse invasion route included *HOPX*, *CKB*, *RFX4*, and *OLIG1*. *HOPX*, a homeodomain-containing transcription factor, is involved in stem cell maintenance [29]. *CKB*, an enzyme, regulates cellular energy homeostasis and is linked to cancer [30]. *RFX4*, a transcription factor recently identified as sufficient for the directed differentiation of CNS cell types from embryonic stem cells [31], *OLIG1*, essential for oligodendrocyte differentiation, contributes to central nervous system myelination [32]. Finally, for the leptomeningeal invasion route, *HMGA1* and *PRRX1* appeared as selective regulators. *HMGA1*, a chromatin-binding protein, is implicated in transcriptional regulation and cancer progression [33]. *PRRX1*, a transcription factor, contributes to embryonic development and cancer invasiveness [34]. *IFI16* and *HBGEF* are growth-condition selective regulators, suggesting that these genes might be explored as markers of GBM cells responding to the tumor microenvironment in future independent work. Intriguingly, the transcription factor *MITF* and some of its known targets (*DCT, MLANA, PLT1* and *S100A1*) - genes implicated in melanogenesis - were detected as a module active in bulk and perivascular invading cells. Moreover, *JUND*, *PDGFRA*, and *SCX* exhibited a high degree of patient selectivity, suggesting that these genes might have applications as robust biomarkers of inter-tumoral variation.

In summary, scregclust identified a concise set of genes with a possible upstream role in determining cell states associated with GBM invasion. To substantiate our findings, we compiled a shorter list of promising regulators to move forward with and validate at the protein level, as discussed next.

### Multispectral protein detection confirms markers of GBM route-specific invasion

Our next objective was to validate the candidate regulator genes at the protein level by assessing their expression in different regions of the mouse GBM xenografts. For this, we combined 6-plex multi immunofluorescence staining with computational image segmentation to measure the expression of each protein in different spatial contexts (Figure 4A, Supplementary file 3, supplementary figure 5). To identify such contexts, we co-stained each protein of interest with markers for tumor cells (anti-human STEM121/NCL), blood vessels (CD31), and white matter (MBP). Utilizing morphological criteria and image k-means clustering, we segmented each slide into 9 different spatial compartments: high-density tumor (1), medium-density tumor (2), low-density tumor (3), circle-shaped aggregates (4), tumor cells growing in close proximity to vasculature (5), diffusively growing cells in the corpus callosum (6), diffusely-growing elongated tumor cells (7), mouse endothelial cells (8) and the mouse brain parenchyma (9) (Figure 4B). As positive controls, we observed that CD31 produced a selective signal in the vascular spatial compartment (number 8), whereas STEM121/NCL was selective for all tumor-containing spatial compartments (Figure 4C). Furthermore, in support of our computational segmentation, we confirmed that it accurately scored the relative abundance of different spatial compartments (e.g., the amount of dense tumor or perivascular cells), consistent with the manually observed phenotype of each PDX model, grouping the cultures into dense/perivascular and diffuse clusters, respectively (Figure 4D).

**Figure 4:**
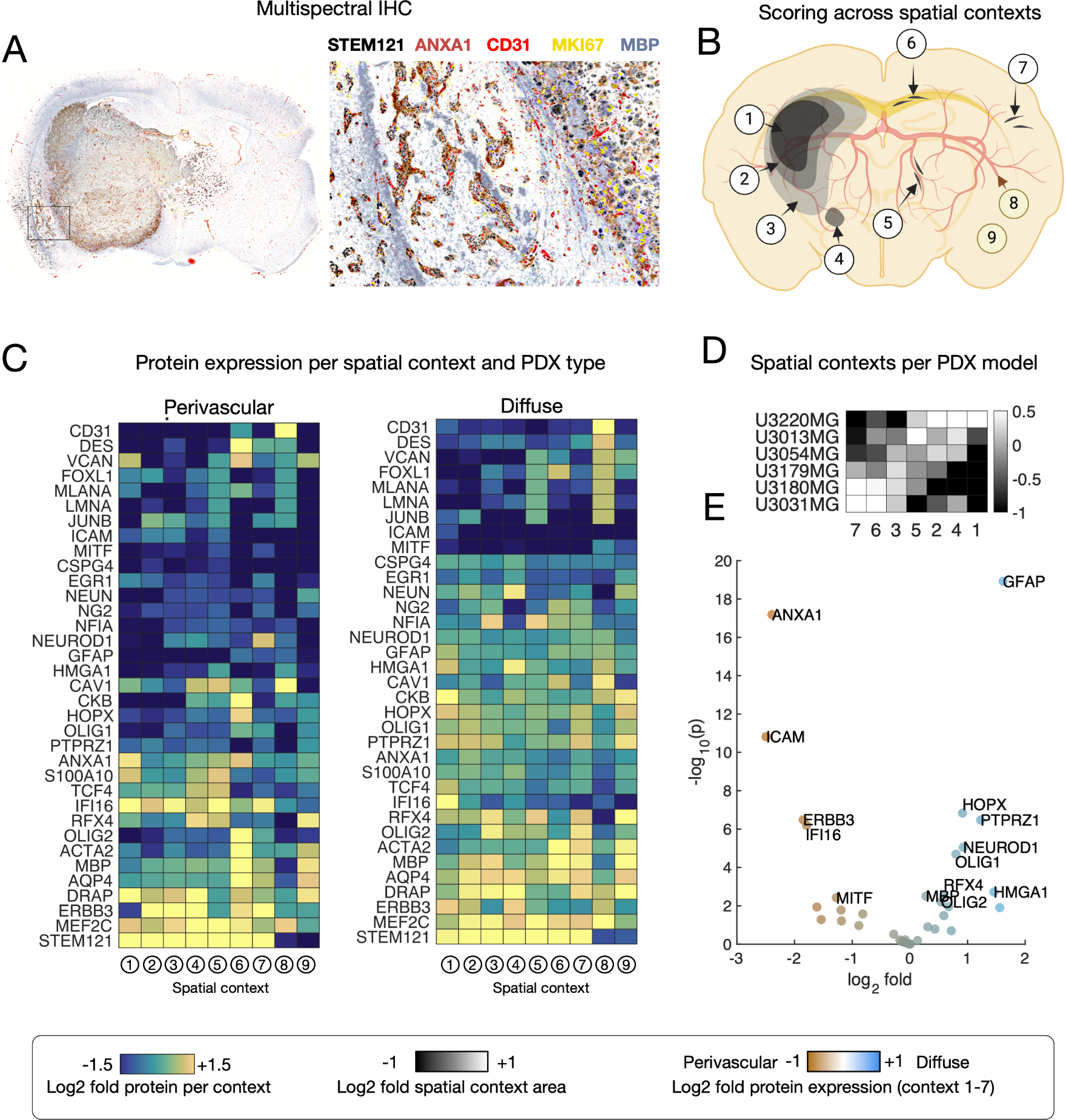
Spatial proteomics reveals route-specific GBM invasion markers. (A) Multispectral IHC of U3054MG PDX, with example staining. (B) We segmented scans into 8 compartments (high-density tumor (1), medium-density tumor (2), low-density tumor (3), circle-shaped aggregates (4), tumor cells growing within close proximity to the vasculature (5), diffusely-growing elongated tumor cells in the corpus callosum (6), other diffusely invading cells (7), blood vessels (8), and mouse brain parenchyma (9). (C) Scoring all PDX models using 35 antibodies; upregulated and downregulated expression of proteins in named compartments for perivascular and diffusively invading cells. (D) Relative area of segmented compartments per PDX cell line. (E) Volcano plot indicating key differentially expressed proteins.

Analysis of all 6 PDX models (n=240 scans) showed that perivascular invading GBM cells exhibited higher expression of ANXA1 and CAV1 protein in their perivascular compartments (numbers 4 and 5) compared to diffusely invading cells (Figure 4E). In line with the neurodevelopmental transcriptional phenotype observed for the diffusely invading cell lines, they displayed a relatively higher abundance of RFX4, AQP4, and HOPX (number 7). Notably, OLIG2 protein was enriched in elongated cells in sparse areas of the tumor, identifying it as a marker of cells that individually penetrate the brain parenchyma. (Figure 4C)

Overall, these findings underscore the heterogeneity of protein expression in invasive GBM and provide further support for ANXA1 protein as a selective marker of perivascular invasion GBM and HOPX and RFX4 as candidate protein markers for diffuse route-invading GBM.

### Validation of *ANXA1*, *HOPX*, and *RFX4* as biomarkers of GBM invasion in patient samples

To assess the translational value of our laboratory findings, we investigated potential regulator expressions in human tissue microarray (TMA) samples from the HGCC biobank (n=148). Given the strong correlation of invasion routes with *ANXA1*, *HOPX*, and *RFX4*, these markers were chosen. Samples showed high expression of *ANXA1* in cells surrounding blood vessels, whereas cells with *HOPX* expression were found away from the blood vessels, which is in accordance with our PDX data (Figure 5A). *RFX4* expression was present in both normal brain tissue and the tumor core area. Next, we questioned whether these markers were associated with patient survival. Interestingly, survival data from HGCC revealed that high *ANXA1*, *HOPX*, and *RFX4* expression in TMA correlate with shorter survival (Figure 5B).

**Figure 5:**
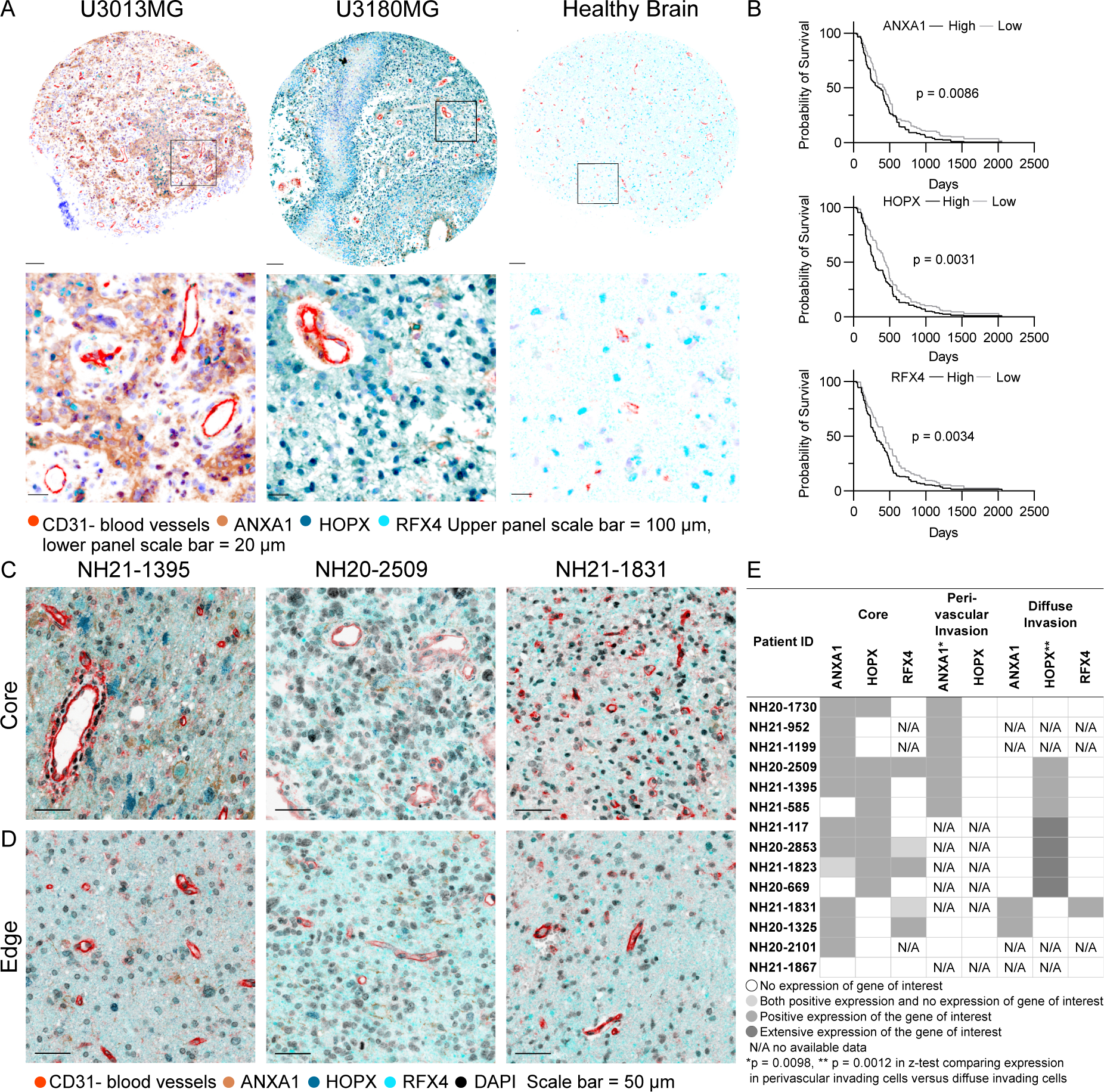
Validation of route-specific invasion markers in an independent patient cohort. (A) Human tissue microarray (TMA) staining of the tumor core, including patients U3013MG, U3180MG, and healthy brain tissue from HGCC. Staining includes CD31 in red, ANXA1 in brown, HOPX in blue, and RFX4 in cyan. The upper panel scale bar indicates 100 μm, while the lower panel scale bar is 20 μm. (B) Survival data from HGCC, showing that high expression of ANXA1, HOPX, and RFX4 correlates with shorter median survival. (C) Staining of the tumor core and (D) edge from three patients from the Queen Square/NHNN repository. Staining includes CD31 in red, ANXA1 in brown, HOPX in blue, RFX4 in cyan, and DAPI in black. The scale bar indicates 50 μm. (E) ANXA1 (p = 0.0098) and HOPX proteins are selectively found in perivascular and diffuse regions in BrainUK samples (N/A means that this type of invasion was absent in the sample).

Although extensive, the HGCC biobank consists of samples of mostly European ancestry patients from a single hospital, and the tumor samples are from an unannotated core region. Therefore, to avoid bias from a single cohort, we also investigated patient tumor samples from an independent cohort, the Queen Square/NHNN repository (ethical approval was obtained via BrainUK, ref:21/014). Also, in this cohort, ANXA1 expression was observed localized to tumor cells near blood vessels both within the tumor core and outside the tumor bulk. Since HOPX is also expressed in normal brain tissue, the distinction of its expression in the tumor core or border region is more challenging. Nevertheless, HOPX was expressed in neurons and glial cells, reflecting our PDX findings. RFX4 expression was found in normal healthy brain tissue and scattered in the tumor core in some patient cases (Figure 5C, D). Importantly, the expressions of ANXA1 and HOPX were found to be invasion-route specific and not patient-specific also within this cohort (Figure 5E).

*ANXA1* has been investigated before in different cancer types [28]. In gliomas, *ANXA1* has been shown to play a role in glioma progression [35], to be present in the immune microenvironment and to be correlated with survival and metastasis potential [36]. Less, however, is known about this protein’s role in perivascular invasion in GBM. *HOPX* plays a critical role during normal development and is strongly expressed in radial astrocyte stem cells [37] and outer radial glial-like cells [25]. *RFX4* functions as a transcription factor and may serve as a potential marker of GBM stem cells [38], with increased expression observed in gliomas [39] and implicated in astrocyte differentiation in cell models [40, 41]. Furthermore, it correlated with poor GBM prognosis [38]. Our confirmation of these markers in two independent patient sample cohorts underscores their value in delineating cell populations potentially driving distinct types of invasion in GBM.

### Mice xenotransplanted with *ANXA1*-KO U3013MG, *HOPX*-KO U3180MG, or *RFX4*-KO U3180MG show increased survival and exhibit a shift in invasion phenotype

Next, we sought to evaluate the impact of *ANXA1*, *HOPX*, and *RFX4* on GBM cell invasion and survival. *ANXA1*, the predicted regulator of perivascular invasion, and *HOPX* and *RFX4*, the predicted regulators of diffuse invasion, were knocked out (KO) with CRISPR/Cas9 in U3013MG and U3180MG, respectively. These two cell lines were chosen due to their capacity for lentiviral modification. Cells were transduced with scramble (SCR) guide RNAs as controls. The KO was confirmed by PCR and sequencing of the flanked region, and by loss of protein expression for the markers expressed *in vitro*. Cell identity was confirmed with STR profiling (Supplementary file 4, and Supplementary figure 6, 7). Before injecting the cells into mice, we conducted proliferation and self-renewal assays *in vitro* to ensure that *ANXA1*-KO, *HOPX*-KO, and *RFX4*-KO cells exhibited no discernible advantages in growth or tumor-forming capabilities (Supplementary figure 8).

We orthotopically injected nude mice with *ANXA1*-KO U3013MG cells, *HOPX*-KO U3180MG cells, and *RFX4*-KO U3180MG cells, along with corresponding SCR control. We then assessed survival, pathology, and gene and protein expression changes.

Mice grafted with *ANXA1*-KO U3013MG cells showed significantly extended median survival time (Figure 6A, p-value < 0,0001) as compared to SCR control. To further analyze the impact of *ANXA1*-KO, we evaluated the brain of the xenografted mice histologically. We observed that *ANXA1*-KO U3013MG tumors did not form a bulk tumor as SCR-U3013MG and U3013MG-WT did (Figure 6D, E). Specifically, the high-density tumor (1), medium-density tumor (2) areas abundance was decreased in *ANXA1*-KO, as well as a decrease of tumor cells growing within close proximity of the vasculature (5). *ANXA1*-KO cells had a higher tendency to grow as a low-density tumor (3), and their morphology shifted from cell aggregates (4) to more diffusely-growing elongated tumor cells (7) (Figure 6D, E, I). In summary, the absence of ANXA1 in tumor cells reduced the density of the tumor bulk, eliminated perivascular invasion and instead shifted the main invasion mode to diffuse invasion. We did not observe significant changes in the number of proliferating cells compared to SCR controls (Figure 6L).

**Figure 6:**
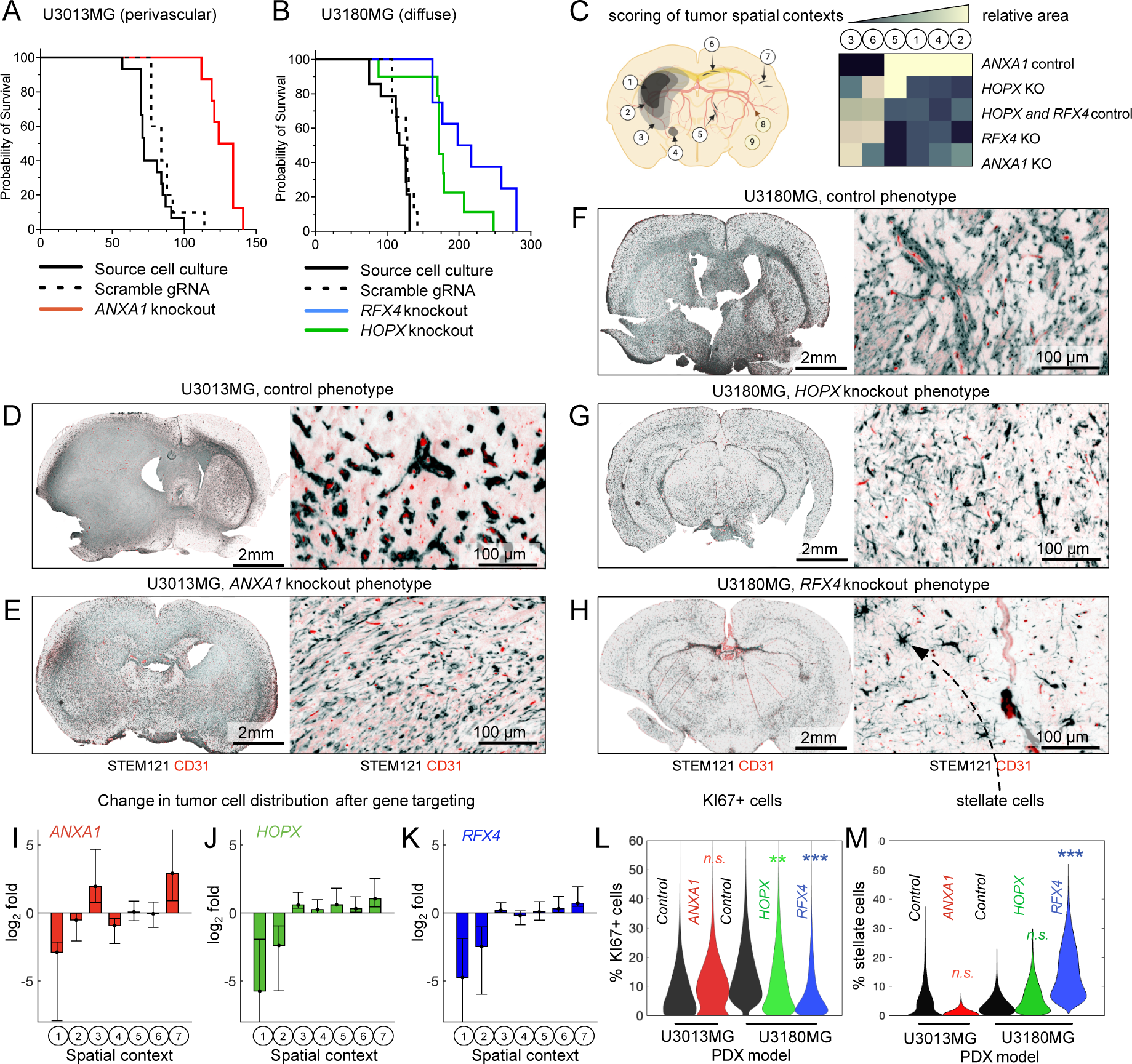
Mouse xenotransplants of *ANXA1*-KO U3013MG, *HOPX*-KO U3180MG, and *RFX4*-KO U3180MG demonstrate prolonged survival and alteration in invasion phenotype. (A,B) Mouse survival for ANXA1-KO U3013MG, *HOPX*-KO U3180MG, and *RFX4*-KO U3180MG. (C) Automated segmentation into 8 compartments. (D-H) Whole brain scans and staining for each geno-type. (I-K) Change in compartment area for each KO-PDX compared to wild-type. (L) Percentage of KI67+ cells in each genotype. (M) Percentage of stellate cells in each genotype.

In the diffusely growing U3180MG xenografts, targeting of either *HOPX* or *RFX4* prolonged survival, decreased tumor cell density, and (in the case of *RFX4*) led to altered morphology of the tumor cells. The KO of *HOPX* in U3180MG increased median survival (Figure 6B, p-value = 0,0002) and these tumors appeared less aggressive than the control, as judged by reduction of tumor density (Figure 6J). We saw no obvious phenotypic change of the tumor cells, except a possible increase in individual tumor cells making contact with blood vessels (Figure 6G, J). The KO of *RFX4* also increased median survival time significantly (Figure 6B, p-value < 0,0001). The *RFX4*-KO PDX showed a marked reduction of tumor cells density (Figure 6H, K). Additionally, a notable number of invading cells, often seen in the striatum, had a stellate phenotype, reminiscent of lower grade glioma (Figure 6H, M).

In summary, these findings confirm *ANXA1*’s role in perivascular invasion in U3013MG grafts and highlight the importance of *HOPX* and *RFX4* in the diffuse growth of U3180MG tumor grafts.

### The shift in preferred invasion route is accompanied by changes in transcriptional cell state

To understand the mechanisms underlying the transition in invasion routes following KO interventions, we performed single-cell profiling of *ANXA1*-KO and *RFX4*-KO tumor cells extracted from mouse brains. Cells from the diffusively invading *ANXA1*-KO tumors exhibited a transition from the MES- and OPC-like states observed in control *ANXA1*-WT tumor cells, favoring NPC- and AC-like states (Figure 7A, B). This trend toward astrocytic differentiation was further supported by differential expression analysis and gene set enrichment analysis, demonstrating a significant upregulation of astrocytic marker *GFAP* (Figure 7C and Supplementary figure 9). Additionally, we observed an upregulation of *GAP43*, a marker of regenerating neurons and reactive glial cells suggested to play a role in GBM invasion [42]. Anecdotally, the transcription factor *MITF* and some of its known targets (*DCT, MLANA, PLT1* and *S100A1*) - genes implicated in melanogenesis - were down-regulated upon *ANXA1* loss (Supplementary figure 9).

**Figure 7:**
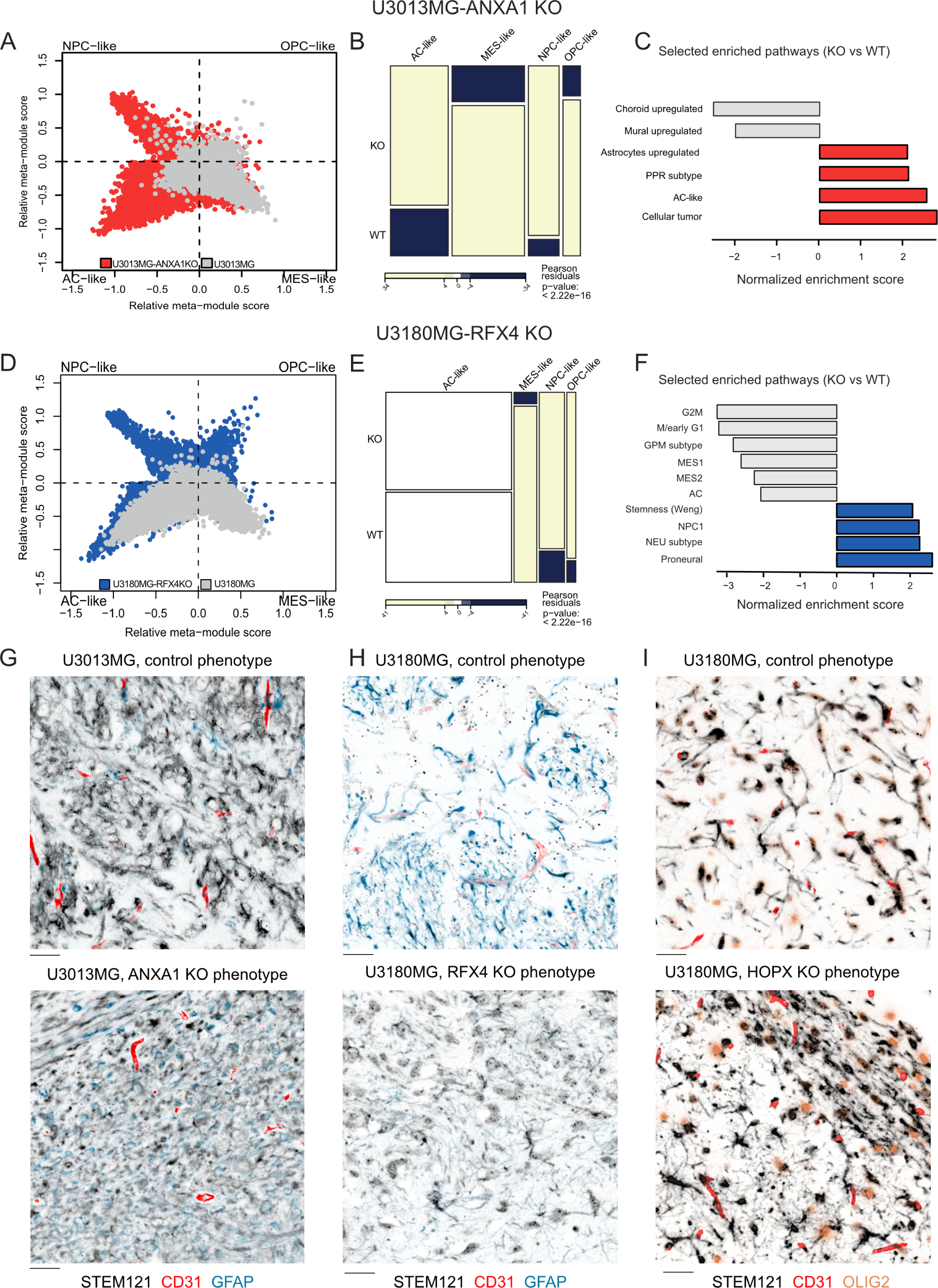
Ablation of *ANXA1*, *RFX4*, and *HOPX* alter GBM cell state distribution and differentiation. (A,B,C) Comparison of *ANXA1* KO U3013MG cells vs wild-type, showing a shift towards NPC-like and AC-like differentiation. (D,E,F) Shift of cell state distribution in *RFX4* KO U3180MG cells, towards an NPC-like, low-proliferating state. (G, H, I) Multispectral IHC staining of (G) GFAP in *ANXA1*-KO U3013MG and U3013MG CTRL, (H) GFAP in *RFX4*-KO U3180MG, and (H) OLIG2 in *HOPX*-KO U3180MG. Scalebar in G, H, I = 25 um

Conversely, in U3180MG xenograft tumors, *RFX4*-KO triggered a significant increase in NPC- and OPC-like cells compared to controls, accompanied by a positive enrichment of gene signatures related to the neuronal lineage (Figure 7D, E, F). Furthermore, the knockout of *RFX4* led to a drastic decrease in the proportion of MES-like cells, along with a negative enrichment of cell cycle-related gene sets. This reduction in actively cycling cells was consistent with the lower tumor cell density in *RFX4*-KO tumors (Figure 6K). Interestingly, the most downregulated mRNA transcript in the *RFX4*-KO samples was *HOPX*, suggesting a regulatory interdependency between the two (Supplementary figure 9).

To validate these findings at the protein level, we conducted additional 6-plex multi immunofluorescence. *ANXA1*-KO tumors revealed a complete abolishment of ANXA1 protein expression, accompanied by increased expression of the astrocyte lineage marker GFAP in tumor cells compared to control (U3013MG SCR) brains (Figure 7G, Supplementary figure 10, p = 0.0000043). In further agreement with the observed alterations in astrocytic signatures, *RFX4*-KO tumors showed decreased expression of GFAP in tumor cells as compared to control (U3180MG SCR) brains (Figure 7H, Supplementary figure 10, p = 0.0014). *HOPX*-KO tumors revealed an increase of OLIG2 expression in tumor cells compared to control (U3180MG SCR) brains (Figure 7I, Supplementary figure 10, p = 0.00023).

In summary, knocking out *ANXA1* led to a transition to a diffuse invasion mode, accompanied by a shift toward an astrocyte-like transcriptional and protein expression profile. Conversely, while *RFX4*-KO and *HOPX*-KO cells still exhibited diffuse invasion, their inherent transcriptional profiles, fraction of cycling cells, and expression of astrocytic and oligodendrocytic protein markers underwent notable alterations.

## Discussion

Extensive invasion, a hallmark characteristic of GBM, contributes to poor prognosis, patient mortality, and relapse. While various invasion routes exist, such as perivascular, diffuse infiltration, or perineuronal satellitosis, the underlying mechanisms remain elusive. Our results provide several new pieces to the puzzle of brain tumor progression. Upon injecting patient-derived cells into the mouse brain, distinct known invasion patterns emerge that correlate with specific transcriptional states: MES-like cells exhibit perivascular invasion, while AC-like and NPC-like cells display diffuse invasion. Using a data-driven modeling strategy, we predicted possible regulators of these states, which were validated in patient samples, *in vivo* experiments, and in-depth molecular profiling.

In this study, *ANXA1* emerged as a pivotal gene in GBM perivascular invasion. Knocking out *ANXA1* in perivascular invading cells induced notable phenotypic shifts, including the loss of tumor bulk and perivascular invasion, while acquiring an AC-like cell state and diffuse invasion, ultimately leading to increased median survival in mice. This observation suggests that the ANXA1+ perivascular invading phenotype potentially drives a reactive cell state, possibly linked to genes associated with injury response [43]. Mechanistically, ANXA1 plays roles in inflammation and tumor cell migration [44]. Cleavage of ANXA1 protein at the cell membrane generates a ligand for formyl peptide receptors, a class of G protein-coupled receptors involved in cell movement. Notably, targeting *ANXA1* increases radiosensitivity in GBM cell lines [35] and is expressed downstream of the ephrin B2 receptor (EFBN2), which is implicated in mouse (G26) models of perivascular invasion [14]. In our data, EFNB2 was not selected by scregclust as a regulator because of its low expression in the human-derived PDX models. The microenvironment, particularly the perivascular niche, significantly influences phenotypic expression, augmenting the perivascular invading phenotype and enriching proteins linked to mesenchymal transformation [45, 46]. Our results appear consistent with the oncostreams phenotype [47], described as the collective invasion of COL1A1-positive tumor cells with mesenchymal properties. We propose that targeting ANXA1 may offer a strategy to suppress COL1A1-positive oncostreams in GBM, possibly with enhanced selectivity compared to targeting collagen 1 directly (since ANXA1 is less abundant in the normal brain than COL1A1). In epithelial cancers, epithelial-to-mesenchymal transition (EMT) has been linked to growth along blood vessels [48]. In accordance, we found intriguing overlaps between previously described regulators of EMT and genes detected in ANXA1-positive cells, including TCF4 [45, 46], and S100A10 [49]. Future research endeavors include delineating regulatory dependencies and exploring the efficacy of targeting ANXA1 with small peptides [50] or exploiting it as a surface antigen for perivascular invading GBM cells. The association of NPC-like and AC-like cells to diffuse growth found in this work aligns well with the phenotype of non-malignant NPCs and astrocytes. Astrocytic and neural precursor migration is an integral part of brain development and injury response [51]. In response to injury, astrocytes transition from a quiescent to a migratory state, contributing to tissue repair and neuronal survival [43].

In contrast to the absolute loss of perivascular invasion upon *ANXA1* ablation, targeting the predicted diffuse invasion drivers *HOPX* and *RFX4* did not result in a complete loss of the phenotype in question, but rather a more complex shift in cell state linked to reduced proliferation and extended median survival in mice. The difference between the intervention experiments may point to the AC as a more robust cell state, consistent with what Schmitt et al. observed, that MES-like cells are more sensitive to reprogramming cues than other GBM states, which are more ‘hardwired’ [52]. Knockout of *RFX4* suppressed AC-like transcriptional signatures and protein expression of GFAP, together with a higher expression of NPC-like signatures. *RFX4* drives the maturation of neural stem cells and neural structures [53, 54], and our results point to a possible role in promoting AC-like states and growth in GBM. The presence of stellate cells in the *RFX4* knockout brains is intriguing and may point to a particular subpopulation that will require further investigation. Our knockout results point to *HOPX* as being the most down-regulated gene upon *RFX4* targeting, potentially suggesting partially a shared mechanism between these two interventions on GBM invasion and growth.

Each of the *ANXA1, RFX4* and *HOPX*-KOs extended mouse survival times. We propose that the extended survival observed in mice grafted with *ANXA1*-KO cells is attributable to the absence of tumor bulk growth and perivascular invasion and the subsequent shift towards diffuse invasion. In these mice, the tumor cells appear more integrated into the brain tissue without forming a bulky mass that exerts pressure. As for the mice lacking HOPX and RFX4, although the exact mechanism is less clear, it’s probable that the reduction in actively cycling cells contributes to their increased survival. While it’s premature to extrapolate these findings directly to human patients, the correlation with improved survival outcomes suggests a potential clinical relevance.

Our results extend and complement previous studies aimed at relating cell differentiation to invasive growth in GBM. Firstly, Brooks et al. proposed a model wherein oligodendrocytic differentiation, dependent on *SOX10*, is observed among cells invading axonally in white matter tracts [22]. While the white matter invading phenotype was not a main focus of this study, our scregclust analysis detected a cluster expressing oligodendrocytic markers, present in two of the cell cultures that we have characterized as bulky and perivascular. While oligodendrocytic protein markers were expressed in these PDXs, we did not, however, see a specific expression of these markers in white matter-located cells (Supplementary figure 11). Secondly, Venkataramani et al. suggested that diffuse invasion is primarily driven by OPC-like and NPC-like cells [18]. Further examination revealed that a significant portion of diffusively invading unconnected cells consists of AC-like and NPC-like cells, supporting our observation that these cells utilize a diffuse invasion route. Lastly, Varn et al. identified two distinct GBM recurrence phenotypes: one characterized as neuronal and the other as mesenchymal, both linked with invasiveness [17]. This further supports both a mesenchymal mode of invasion and a neuronal mode of migration for the invasive GBM cells remaining in the normal brain parenchyma. Further work is needed to refine the nomenclature around cell states and invasion routes in GBM, and the association between AC-like cells and diffuse growth consistently observed across our three diffusely growing PDX models extends previous observations. Towards this goal, studying GBM invasion across a larger clinical repertoire will be crucial. This would, for instance, open for robust statistical associations between tumor genetic and epigenomic features and their morphological presentation.

Methodologically, our study introduces a framework for uncovering invasive cell states and their regulators. In this study, we employ scregclust to identify key gene regulators implicated in perivascular or diffuse invasion, leveraging scRNA-seq data. Subsequently, we validate the protein expression of these regulators within the invasion niche of patient samples from two independent cohorts using multiplex immunofluorescence staining. Upon perturbation of a potential regulator in the invasion route of interest, we observe significant alterations in both RNA and protein expression, impacting the invasion route, migratory behavior, and morphology of these cells *in vivo*. Just like the observed transcriptional states are much more pronounced in the brain environment compared to adherent cultures, the effect of gene targeting is more pronounced *in vivo* than *in vitro*. It thus appears crucial to anchor the discovery of regulators of invasion in sufficiently complex models that recapitulate at least crucial parts of the brain environment. We acknowledge that our immunodeficient mouse models lack central aspects, which makes it important to validate the discovered functional biomarkers in independent patient materials, as was done here.

Taken together, this work presents a scalable approach to uncover critical genes that underlie specific cell states linked to brain tumor invasion. Looking ahead, it will be important to extend investigations to larger clinical repertoires, and to leverage our understanding of invasion regulators to interfere with these processes in a tailored manner. We reserve this for future work.

## Acknowledgements

We thank the involved patients and their families for the support and donation of materials to the Human Glioblastoma Cell Culture (HGCC) biobank. We also thank the HGCC team for invaluable contributions in collecting and providing the patient derived cell cultures used in this study. We thank the ongoing HGCC Tissue Microarray (Tobias Bergström, unpublished) and HGCC Phenobank initiative (Cecilia Krona, unpublished) for sharing TMAs and phenotypic data, respectively. We thank the Brain UK biobank for making patient materials investigated in Figure 5 available. We thank the National Genomics Infrastructure (NGI) for providing the sequencing service and the BioVis Platform for providing assistance with FACS sorting and microscopy. We thank FoUU for assistance with sectioning and scanning tissue slides and Artur Mezheyeuski for sharing expertise on multiplex staining, multispectral image acquisition, and data analysis.

## Author contributions

Experiments were performed by MD, RS, IY, MBB, JH, LE, and RE. Mouse experiments were performed by SK, RS, IY, MBB, MD, and CK (coordination). Profiling and imaging data were analyzed by MD, IL, and SN. Human tissue was analyzed by TM and SM. The first manuscript draft was prepared by RS, IY, IL, SK, MBB, MD, SN, with input from the other authors. SN guided the study.

## Declaration of generative AI and AI-assisted technologies in the writing process

The spelling and grammar of the manuscript were checked using Grammarly (Grammarly, Inc.) and ChatGPT-3.5 (openAI, Inc.). All text was reviewed and edited by the authors, and the authors take full responsibility for the content written for the publication.

## Methods

### 1. Patient Samples

Patient-derived glioblastoma cell lines were established from tumor tissue as previously explained (Reference [23]). All samples were collected with the informed consent of the patients, and the collection was approved by the Uppsala Regional Ethical Board, under number 2007/353. The cells were seeded on 1% laminin-coated flasks and maintained in serum-free neural stem cell medium with B-27 and N2 supplements, as well as EGF and FGF growth factors. The experiments adhered to the principles outlined in the WMA Declaration of Helsinki and the Department of Health and Human Services Belmont Report.

### 2. Mouse Xenografts: Patient-derived xenograft model

All mouse experiments were conducted in strict accordance with an ethical permit granted by the Uppsala Animal Research Ethical Board, bearing reference numbers C41/14 and 5.8.18-06726/2020. NMRI nude (NMRI-Foxn1 nu/nu) mice were procured from Janvier Labs, while Hsd:Athymic nude-Foxn1 mice were procured from Envigo. Mice falling within the age range of 6 to 9 weeks were selected for the experiments. They were housed in individually ventilated cages, with each cage accommodating up to 5 mice. Appropriate housing enrichment, bedding material, food, and drinking water were provided ad libitum, and the mice were maintained on a 12/12-hour light cycle. Human glioma cell cultures demonstrating verified tumor growth and the desired phenotype were systematically labeled with a lentivirus expressing GFP-Luciferase to enable subsequent tracking. All cell lines were STR profiled before injections to confirm their genetic identity (Eurofins Genomics). Orthotopic tumor injections were carried out by transplanting 100,000 labeled cells into the striatum of each mouse. The mice were monitored using luciferase imaging, weight measurements, and onset of neurological symptoms for up to 40 weeks post-injection. Upon reaching the defined experimental endpoints, the brains of the terminal mice were harvested and processed for histology analyses or cell isolation.

### 3. Immunohistochemistry

For histology, mouse brains were processed in an automated tissue processor (TPC) under the following conditions: 1 hour 70% EtOH, 2x 1 hour 96% EtOH, 3x 1 hour 100% EtOH, 2x 1 hour Xylene, and 3x 1 hour Paraffin at 60°C. The paraffin-embedded brains were then sectioned into 5µm slides. Each block was analyzed for protein expression using a standard IHC protocol. In brief, after deparaffinization, antigen retrieval using Antigen Unmasking Solution Citrate-Based pH 6 (Vector Laboratories #H-3300) with 0.05% Tween 20 (Biorad #1610781) was commenced in 2100 Antigen Retriever for 15 min followed by cooling down to room temperature. Then sections were incubated in 3% H2O2 (30% H2O2 (Thermo Scientific #10687022) diluted in TBS) for 10 min, followed by washes with TBS-T (washing buffer). The sections were blocked with Normal Antibody Diluent (ImmunoLogic WellMed #UD09) for 30 min at room temperature, and primary antibodies STEM121 (1:500) (Takara #Y40410), ANXA1 (1:400), RFX4 (1:500) (HPA #050527), and HOPX (1:1000) diluted in Normal Antibody Diluent were applied and incubated for 60 min at RT. We used BrightVision, a 1-step detection system, Goat Anti-Rabbit HRP (WellMed #DPVR110HRP), and anti-Mouse HRP (WellMed #DPVM110HRP) detection systems followed by incubation with Bright-DAB substrate kit (WellMed #KBS04-110). Slides were then counterstained with Myers’ Hematoxylin and permanently mounted with Pertex (HistoLab #00811).

### 4. Multiplex fluorescent staining and multispectral imaging

Multiplex staining was performed using the Opal 6-Plex Manual Detection kit (Akoya Biosciences, NEL861001KT). Procedures were conducted according to the protocol with a deviation, where anti-mouse HRP (Immunologic #DPVM110HRP) and anti-rabbit HRP (Immunologic #DPVR110HRP) were used for antibody detection instead. Antibodies were stripped after each Opal incubation using the microwave, and the procedure was repeated for the next primary antibody-Opal pairing. Every antibody-Opal pairing was independently validated as per the manufacturer’s instructions. The validated antibody-Opal pairings are available in Supplementary file 3. Slides were mounted with ProLong Diamond Antifade Mountant (ThermoFisher #P36970), imaged using the PhenoImager whole slide workflow, and unmixed using InForm 4.8 (Akoya Biosciences) software.

### 5. Single cell isolation from PDX tumors

Upon the experimental endpoint, mouse brains were harvested into cold HBSS buffer containing 1% Pen/Strep, 0.6% glucose, and 25nM HEPES. Then, the brain was sliced using a 1mm coronal section matrix, cut into about 1-2 mm pieces using a surgical blade, and dissociated into single cells using the Tumor Dissociation kit, human (Miltenyi, #130-095-929), used in combination with the Mouse Cell Depletion Kit (Miltenyi, #130-104-694) according to the manufacturer’s protocols. Red blood cells were removed using the Red Blood Cell Lysis Solution (Miltenyi, #130-094-183).

### 6. Single-cell RNA sequencing data generation

The generation of single-cell RNA sequencing libraries followed the manufacturer’s guidelines, utilizing the Chromium Single Cell 3’ Library and Gel Bead Kit v2, v3, and v3.1 (analysis of KO-cells) (10x Genomics, Pleasanton, CA). Cryo-preserved cells underwent washing and re-suspension in 0.1% BSA in PBS just before loading onto a Chromium Single Cell B Chip (10X Genomics) with the aim of capturing 10,000 cells. Subsequently, the quality of the libraries was assessed using Agilent High Sensitivity DNA Kit and Agilent Bioanalyzer 2100 DNA Kit (Agilent Technologies). Libraries were sequenced on an Illumina NovaSeq 6000 with the sequencing configurations recommended by 10X Genomics. Demultiplexing, counting, and alignment to the human (GRCh38) reference genome were performed using Cell Ranger 3.0.2 (10X Genomics).

### 7. Data processing, integration, and cell clustering

We performed single-cell analysis using the Seurat package (v. 4) (Butler et al., 2018). We filtered out cells expressing fewer than 500 genes and genes that were expressed by fewer than 10 cells. We filtered out potential doublets by setting nFeature_RNA parameters at greater than 7200 for v3 of the kit and greater than 5100 for the v2 kit. We removed low-quality cells that contained more than 30% mitochondrial genes, resulting in 110,458 cells retrieved (85.6% of the original population). We also removed highly expressed genes that are not related to the study, such as abundant ribosomal, mitochondrial, and hemoglobin genes. Lastly, we performed cell cycle regression to ameliorate the impact of the cell cycle on cell groupings. Then, we used the reciprocal PCA (RPCA) method to integrate the data and clustered cells using the Louvain algorithm with multilevel refinement. We used a range of resolutions from 0.01 to 1 to unravel cell sub-populations, and based on a dendrogram calculated using the Clustree (v. 0.5.0) package, we assessed the stable and therefore optimal cluster separation.

### 8. Regulatory landscape analysis by scregclust

The scregclust algorithm [26] was applied to the scRNA-seq data from each sample individually. For each run, the initial cluster number was set to 20, a minimum number of genes per cluster to 10, and a range of penalization values were tested (0.01-0.1). The final penalization for each sample was chosen based on the metrics “predictive R2” and “regulator importance,” as described in [26]. For each sample, this resulted in a regulatory table with regulators (transcription factors and kinases) as rows and gene modules as columns. The regulatory tables for all samples were merged into a common table for the entire sample set, and the data were z-transformed (Figure 3A). Modules (columns) were clustered using hierarchical clustering. Gene modules were characterized by quantifying their overlap with gene signatures of GBM cell states and cell cycle phases [8], as well as signatures representing the invasion routes (diffuse, perivascular, leptomeningeal). The overlap was quantified using the Jaccard index. Analysis of variance (ANOVA) tests were performed to assess the specificity of the predicted regulators in regard to growth condition, patient, and invasion route (Figure 3B). Modules were given categorical annotations; the first two were derived from their sample origin (growth condition: *in vitro/in vivo*, and patient: U3013MG, U3031MG, etc.). The third, invasion route, was derived from the above-described scoring. **9. Metamodules.** Metamodules were defined through hierarchical clustering of the merged regulatory table described above, and cutting the dendrogram at height 36, which resulted in 13 metamodules. Signatures were defined by, for each metamodule, merging the gene content of each individual gene module and keeping genes that were common for four gene modules or more. These metamodule signatures were then used to assign a metamodule score to each cell in the dataset provided by [27], using the AddModuleScore()-function in the Seurat R-package.

### 10. Spatial analysis of PDX tumors

To score the intensity of different protein markers in different anatomical niches, we processed the VP images as follows. We regard each pixel as a point in 8-dimensional space (*z*_1_*, z*_2_*, …, z*_8_) where four of the channels were common to all analyzed images: nuclei (DAPI), auto-fluorescence (AF), tumor cells STEM121/NCL, and endothelial cells (CD31). The four remaining channels were used to evaluate proteins of interest. Images were loaded from qptiff format using bioformats toolbox in Matlab. For each channel, we used L2-regularized regression to correct for shared variation with the other channels. The L2 penalty was set to 0 for DAPI and AF channels, and to a tuning constant for the others. After correction, we segmented a four-channel image consisting of the DAPI, AF, STEM121/NCL and CD31 channels using image k means segmentation (Matlab image analysis toolbox), with k set to 5. This consistently resulted in a 5-cluster solution with easily identifiable centroids representing tumor cells (high STEM121/NCL) and endothelial cells (high CD31). Pixels assigned to these centroids were used to obtain binary images T and V representing the tumor (T) and vascular (V) parts of the section. The endothelial niche was defined as the set of positive pixels in the V image. High, medium and low density regions of T were found as positive pixels of the T image in regions of different density, as measured by the Matlab imboxfilter method. We subsequently used a set of morphological property filters to detect elongated tumor cells, tumor cells near blood vessels, tumor cells near vasculature, and tumor cells in dense bundles. After these steps, we had obtained a labeling matrix L, that provided the class of each pixel. We subsequently scored each protein i by measuring its average intensity in each class j, correcting for cellularity using the DAPI channel, i.e.

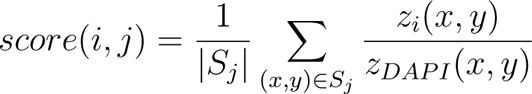

where Sj is the set of pixels (x,y) in class j.

### 11. Tissue microarray study from HGCC cohort and survival analysis

The stitched pyramidal OME-TIFF files were loaded into QuPath 4.3 software [55], and TMA was disarrayed to assign coordinates to the TMA cores. The nuclei were segmented using the StarDist 4.0 extension [56]. Then, for each protein marker, we evaluated staining specificity and set up manual classification thresholds depending on their localization. These fixed thresholds for each marker protein were then used for the classification of all TMAs within the set. The process was separately iterated for each staining set. To see if the expression of protein markers is a hazard factor for patient survival, we performed a survival analysis based on Cox (Proportional Hazards) Regression using the fraction of cells expressing modality in each core (number of cells expressing/total number of nuclei) using the survival (3.4-0) package. For the Kaplan-Meier curves, we stratified the calculated fractions into “high” and “low” based on the median expression value for each marker.

### 12. BrainUK cohort

To extend our collection and provide material that consisted of invasive regions of glioblastoma, we applied for access to samples from BrainUK (BRAIN UK Ref: 21/014). We then checked the expression of our top candidate proteins ANXA1, RFX4, and HOPX using mIF staining as described above. The material was carefully analyzed and scored by neuropathologists in 7-10 fields of view per section, selected in tumor-invaded niches.

### 13. Lentiviral transduction

For generating knockout clones of target genes (*HOPX*, *ANXA1*, *RFX4*, and scramble) for U3013MG and U3180MG, cell cultures were transduced using a reverse-transduction method. Briefly, cells were detached using TrypLe, washed in PBS, and counted. Then, 100,000 cells were co-transduced with the Cas9-nickase and gRNA vectors (see Supplementary file 4). To minimize off-target effects, cells were transduced with the Cas9-nickase vector at MOI 3 and the gRNA vectors at MOI 5. After vector addition, cells were incubated for 2 hours at 37°C, then plated onto laminin-coated 6-well plates. The virus-containing medium was replaced after 24 hours, and selection medium was added 3 days post-transduction. Cultures were treated with antibiotic selection medium for 7-10 days and then passaged for seeding each of them into 96-well plates as single-cell clones using FACS. See Supplementary file 4 for vectors and guideRNA sequences.

### 14. Genotyping PCR and Sanger sequencing

Single-cell clones constituting colonies were genotyped to identify knockout clones. DNA was isolated using lysis buffer and incubated for 2 hours at 60°C. DNA was precipitated using precipitation buffer for 30 minutes at RT and washed 3 times with 70% EtOH. The pellet was air-dried for 30 minutes and then resuspended in TE buffer (pH 8). Clones with visible alterations in amplicon size from high-throughput PCR were selected for Sanger sequencing. In the second step, KAPA HiFi HotStart ReadyMix was used to amplify the DNA, and the amplicons were separated on a 2% agarose gel. The purified amplicons were then subjected to Sanger sequencing. Details of primers used are in Supplementary file 4..

### 15. Knockout evaluation

Sanger sequencing results were qualified and analyzed using Snap-Gene and the ICE CRISPR analysis tool. Clones with knockout indication were expanded, and about 10 million cells were collected from each clone to create FFPE cell pellets for IHC analysis of protein. A small pellet was also collected for second genotyping PCR and sent for Sanger sequencing. FFPE cell pellets were sectioned and stained with antibodies and protocol indicated in Section 2. From clones with confirmed protein loss, a pellet of 100 thousand cells was collected and sent for STR profiling. 3-10 knockout clones per target were mixed in equivalent numbers 6 days before injection in mice.

### 16. Proliferation and self-renewal assays on knockout clones

To assess the proliferation and self-renewal capacities of knockout cells, we used CyQuant Cell Proliferation Assay and Extreme Limiting Dilution Assay (ELDA). In the proliferation test, cells were seeded in a range of serial dilutions in duplicates and allowed to grow for 72 hours. After that, the Cyquant Protocol was performed according to the manufacturer’s instructions. Self-renewal was tested by seeding cells in dilutions ranging from 200 to 1 cell per well in 96-well ultra-low attachment plates over the period of 7 days, two biological replicates were used. ELDA analysis was conducted using software accessible at http://bioinf.wehi.edu.au/software/elda/, following the specified procedure.

### 17. Statistical analyses

#### Cell-state plots

Cell-state plots were generated as described. Barplots in Figure 7B and 7D were generated by counting the number of cells in each quadrant.

#### Mosaic plot

To statistically assess the relation between cell state and invasion route, a chi-square test was performed and visualized as a mosaic plot.

#### Differential gene expression analysis and MA plots

Differential gene expression (DGE) analysis was performed using the FindAllMarkers-function in the Seurat package. MA plots were created by plotting the log2FC-values from the DGE analysis against the log2 total gene count for each gene across cells.

#### Survival analysis (Kaplan-Meier)

Survival data were utilized to create Kaplan-Meier survival curve plots, and the statistical analysis employed the Mantel log-rank test. The median survival is denoted as MS.

## Supplemental information titles and legends

- Supplementary file 1: Phenotypic scores for mice injected with U3013MG, U3054MG, U3220MG, U3179MG, U3180MG, and U3031MG
- Supplementary file 2: SUPPL_S2_scregclust_results
- Supplementary File 3: Vectra Polaris panels, list of antibodies used.
- Supplementary File 4: Constructs used for generating hCas9 nickase and dual guideRNA sequences, Primers used for genotyping and Sanger sequencing.
- Supplementary figure 1: STEM121 IHC for mice injected with U3013MG, U3054MG, U3220MG,
- Supplementary figure 2: STEM121 IHC for mice injected with U3179MG, U3180MG, and U3031MG
- Supplementary figure 3: Separate UMAP and cell state projections of each cell line
- Supplementary figure 4: Metamodules detected by scregclust
- Supplementaryl Figure 5: Segmentation
- Supplementary figure 6: PCR, protein, STR to document the ANXA1 CRISPR-Cas9 knockouts
- Supplementary figure 7: PCR, protein, STR to document the RFX4/HOPX CRISPR-Cas9 knockouts
- Supplementalary figure 8: In vitro characterization of each knockout.
- Supplementary figure 9 (figure): differential expression analysis of the *RFX4*, *HOPX*, and *ANXA1* knockout GBM cultures compared to control.
- Supplementary figure 10 (figure): log2 fold protein expression vs scramble control of STEM121+ cells.
- Supplementary figure 11 (figure): ERBB3 positive U3013MG cells in the corpus callosum.

