## Supplementary Material for "The invasion phenotypes of glioblastoma depend on plastic and reprogrammable cell states"

| Mouse index | Cell culture | Bulk formation | Abnormal blood vessels | Leptoeningeal spread | Diffuse infiltration | Perivascular invasion | White matter invasion | Subpial invasion | Perineural invasion | Mouse survival |
|---|---|---|---|---|---|---|---|---|---|---|
| SSF2_3013 #15 | U3013MG | 1 | 1 | 1 | 0 | 1 | 1 | 0 | 0 | 91 |
| SSF2_3013 #2 | U3013MG | 1 | 0 | 1 | 0 | 1 | 0 | 0 | 0 | 57 |
| SSF2_3013 #3 | U3013MG | 1 | 1 | 1 | 0 | 1 | 1 | 1 | 0 | 71 |
| SSF2_3013 #4 | U3013MG | 1 | 0 | 0 | 0 | 1 | 1 | 1 | 0 | 70 |
| SSF2_3013 #5 | U3013MG | 1 | 0 | 1 | 0 | 1 | 1 | 0 | 0 | 85 |
| SSF2_3013 #6 | U3013MG | 1 | 0 | 1 | 0 | 1 | 0 | 0 | 0 | 71 |
| SSF2_3013 #9 | U3013MG | 1 | 0 | 1 | 0 | 1 | 1 | 0 | 0 | 70 |
| SSF2_3013 #11 | U3013MG | 1 | 0 | 1 | 0 | 1 | 1 | 0 | 0 | 70 |
| SSF2_3013 #12 | U3013MG | 1 | 0 | 1 | 0 | 1 | 1 | 0 | 0 | 87 |
| SSF2_3013 #13 | U3013MG | 0 | 0 | 1 | 0 | 1 | 0 | 0 | 0 | 100 |
| SSF2_3013 #14 | U3013MG | 0 | 0 | 1 | 0 | 1 | 0 | 0 | 0 | 84 |
| SSF2_3031 #5 | U3031MG | 0 | 0 | 1 | 0 | 0 | 1 | 0 | 0 | 106 |
| SSF2_3031 #6 | U3031MG | 1 | 0 | 0 | 0 | 0 | 0 | 0 | 0 | 106 |
| SSF2_3031 #7 | U3031MG | 0 | 0 | 1 | 0 | 0 | 1 | 0 | 0 | 124 |
| SSF2_3031 #8 | U3031MG | 1 | 0 | 1 | 1 | 0 | 1 | 0 | 1 | 124 |
| SSF2_3054R1 #4 | U3054MG | 1 | 1 | 1 | 0 | 1 | 0 | 0 | 0 | 105 |
| SSF2_3054R1 #5 | U3054MG | 1 | 0 | 1 | 0 | 1 | 0 | 0 | 0 | 128 |
| SSF2_3054R1 #8 | U3054MG | 1 | 1 | 1 | 0 | 1 | 0 | 0 | 0 | 134 |
| SSF2_3054R1 #9 | U3054MG | 1 | 0 | 1 | 0 | 1 | 0 | 0 | 0 | 134 |
| SSF2_3054R1 #10 | U3054MG | 0 | 0 | 1 | 0 | 1 | 0 | 0 | 0 | 105 |
| SSF2_3054 #10 | U3054MG | 1 | 1 | 0 | 0 | 1 | 0 | 0 | 0 | 160 |
| SSF2_3054 #12 | U3054MG | 1 | 0 | 0 | 0 | 1 | 0 | 0 | 0 | 113 |
| SSF2_3054 #13 | U3054MG | 1 | 0 | 0 | 0 | 1 | 0 | 0 | 0 | 155 |
| SSF2_3054 #14 | U3054MG | 1 | 0 | 0 | 0 | 1 | 0 | 0 | 0 | 90 |
| SSF2_3054 #1 | U3054MG | 1 | 1 | 1 | 0 | 1 | 0 | 0 | 0 | 155 |
| SSF2_3054 #5 | U3054MG | 1 | 1 | 1 | 0 | 1 | 0 | 0 | 0 | 163 |
| SSF2_3179 #1 | U3179MG | 0 | 0 | 1 | 1 | 1 | 1 | 1 | 1 | 176 |
| SSF2_3179 #3 | U3179MG | 0 | 1 | 1 | 1 | 1 | 1 | 1 | 1 | 210 |
| SSF2_3179 #6 | U3179MG | 1 | 0 | 1 | 1 | 1 | 1 | 1 | 1 | 197 |
| SSF2_3179 #9 | U3179MG | 1 | 0 | 0 | 1 | 0 | 1 | 1 | 1 | 197 |
| SSF2_3180 #2 | U3180MG | 0 | 1 | 0 | 1 | 0 | 1 | 1 | 1 | 114 |
| SSF2_3180 #3 | U3180MG | 0 | 0 | 1 | 1 | 1 | 1 | 1 | 1 | 131 |
| SSF2_3180 #4 | U3180MG | 0 | 0 | 1 | 1 | 0 | 1 | 1 | 1 | 126 |
| SSF2_3180 #5 | U3180MG | 0 | 0 | 1 | 1 | 1 | 1 | 1 | 1 | 126 |
| SSF2_3180 #6 | U3180MG | 0 | 0 | 1 | 1 | 1 | 1 | 1 | 1 | 125 |
| SSF2_3180 #7 | U3180MG | 0 | 0 | 1 | 1 | 0 | 1 | 1 | 1 | 113 |
| SSF2_3180 #9 | U3180MG | 0 | 1 | 1 | 1 | 1 | 1 | 1 | 1 | 117 |
| SSF2_3180 #10 | U3180MG | 0 | 0 | 1 | 1 | 0 | 1 | 1 | 1 | 131 |
| SSF2_3220 #1 | U3220MG | 1 | 1 | 1 | 0 | 1 | 0 | 0 | 0 | 70 |
| SSF2_3220 #2 | U3220MG | 0 | 0 | 0 | 0 | 1 | 1 | 0 | 0 | 58 |
| SSF2_3220 #3 | U3220MG | 0 | 0 | 1 | 0 | 1 | 1 | 0 | 0 | 63 |
| SSF2_3220 #5 | U3220MG | 0 | 0 | 1 | 0 | 1 | 0 | 0 | 0 | 70 |
| SSF2_3220 #6 | U3220MG | 1 | 1 | 1 | 0 | 1 | 1 | 0 | 0 | 70 |
| SSF2_3220 #8 | U3220MG | 1 | 1 | 1 | 0 | 1 | 1 | 0 | 0 | 65 |
| SSF2_3220 #10 | U3220MG | 1 | 0 | 1 | 0 | 1 | 0 | 0 | 0 | 79 |
| average percent agreement | | 72 | 74 | 80 | 96 | 88 | 83 | 97 | 96 | |

(the average fraction of phenotypic observations that are the majority observation in each PDX)

**Supplementary Table 1: GBM PDX models; phenotypes and phenotype consistency.** Presence (1) and absence (0) of different growth phenotypes in mice orthotopically grafted with different primary glioblastoma lines.

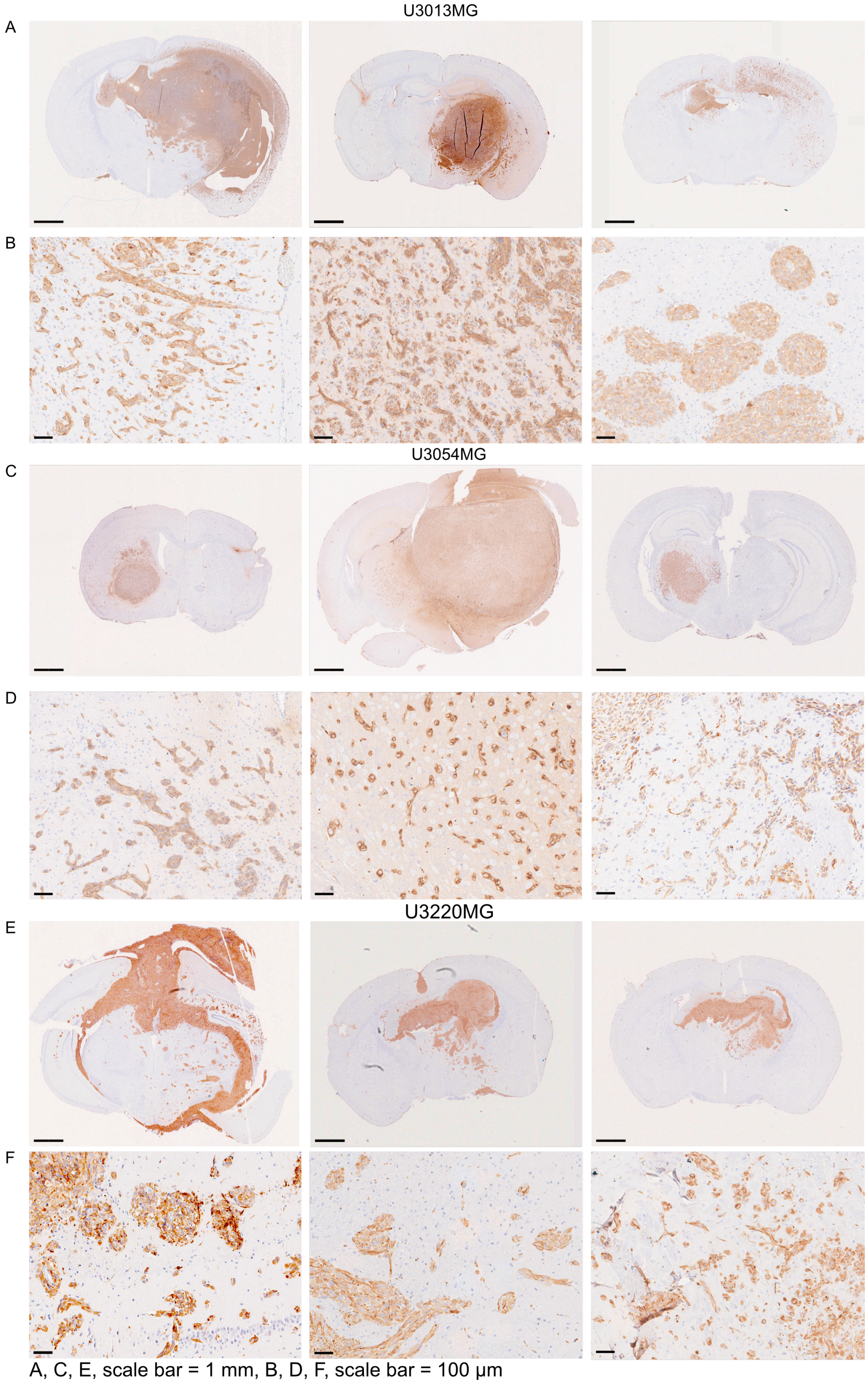


**Supplementary Figure 1: Representative growth phenotypes in mice orthotopically grafted with different primary glioblastoma lines.** (A) U3013MG (B) Detailed view of invasion phenotype of U3013MG (C) 3054MG (D) Detailed view of invasion phenotype of U3054MG (E) 3220MG (F) Detailed view of invasion phenotype of U3220MG. (A, C, E) Scale bar indicating 1mm. (B, D, F) Scale bar indicating 100 µm.

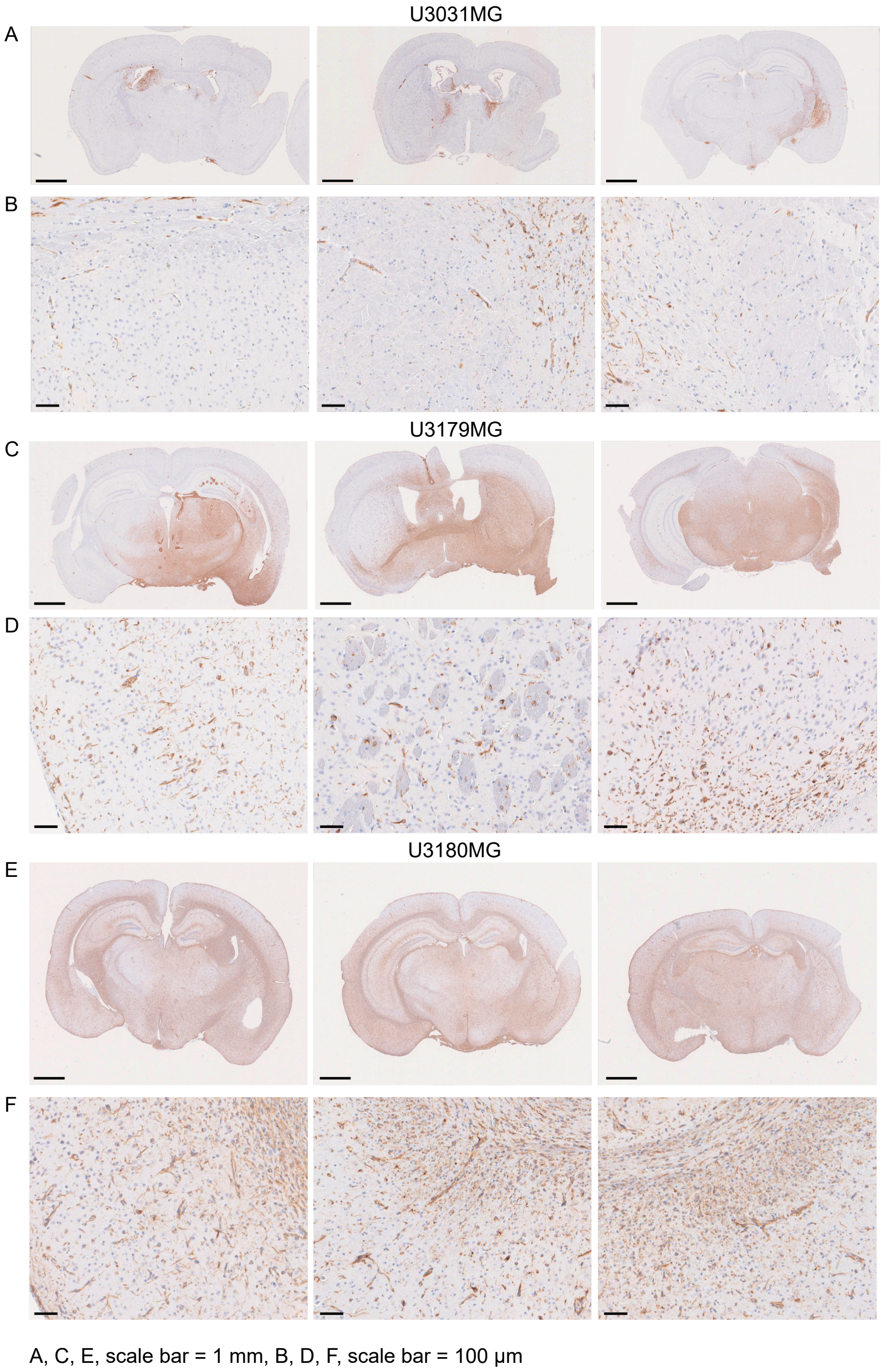


**Supplementary Figure 2: Representative growth phenotypes in mice orthotopically grafted with different primary glioblastoma lines** (A) U3031MG (B) Detailed view of invasion phenotype of U3031MG (C) 3179MG (D) Detailed view of invasion phenotype of U3179MG (E) 3180MG (F) Detailed view of invasion phenotype of U3180MG. (A, C, E) Scale bar indicating 1mm. (B, D, F) Scale bar indicating 100 μm.

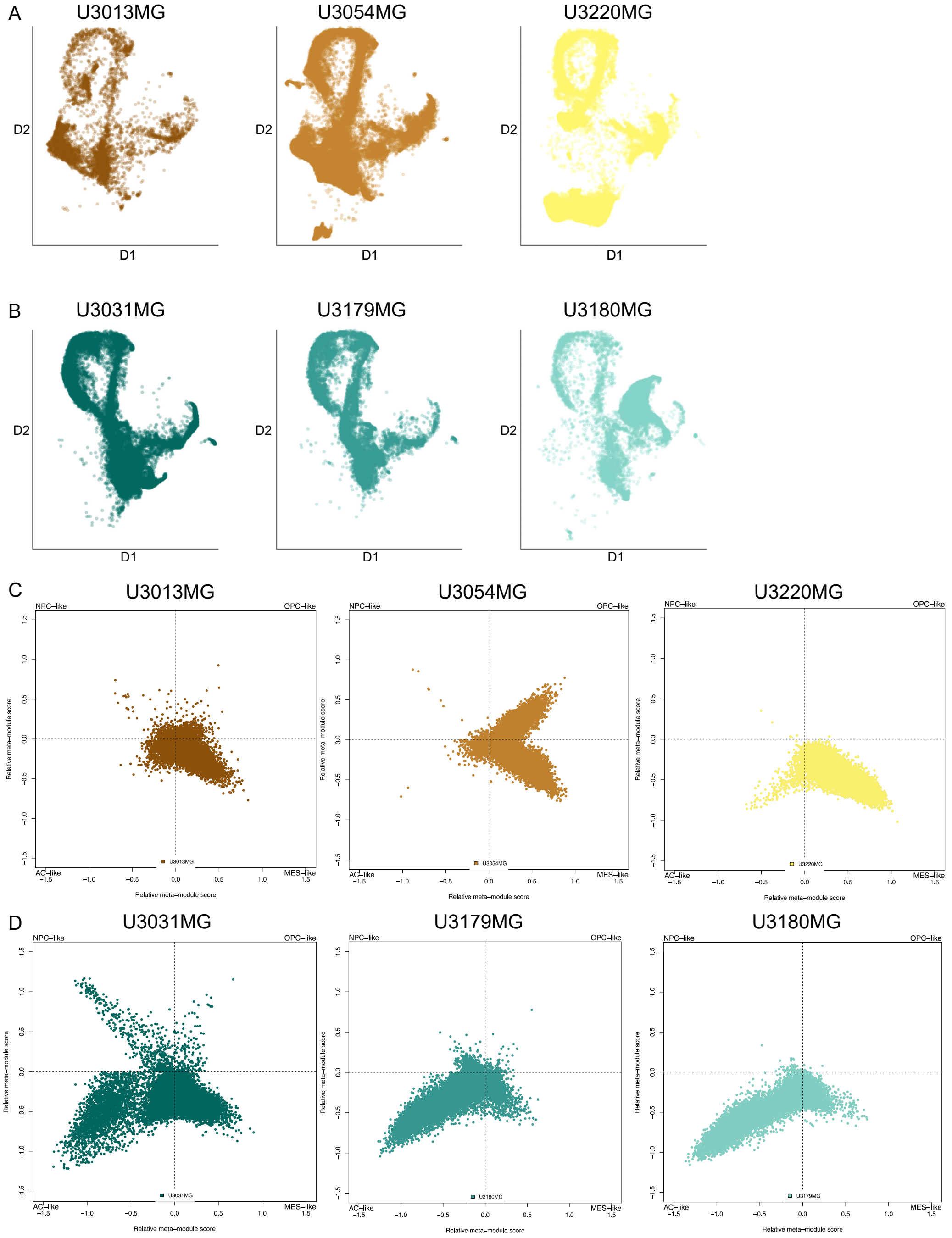


**Supplementary Figure 3: UMAP projections and cell state plots** for each separate sample in Figure 2 of the main manuscript.

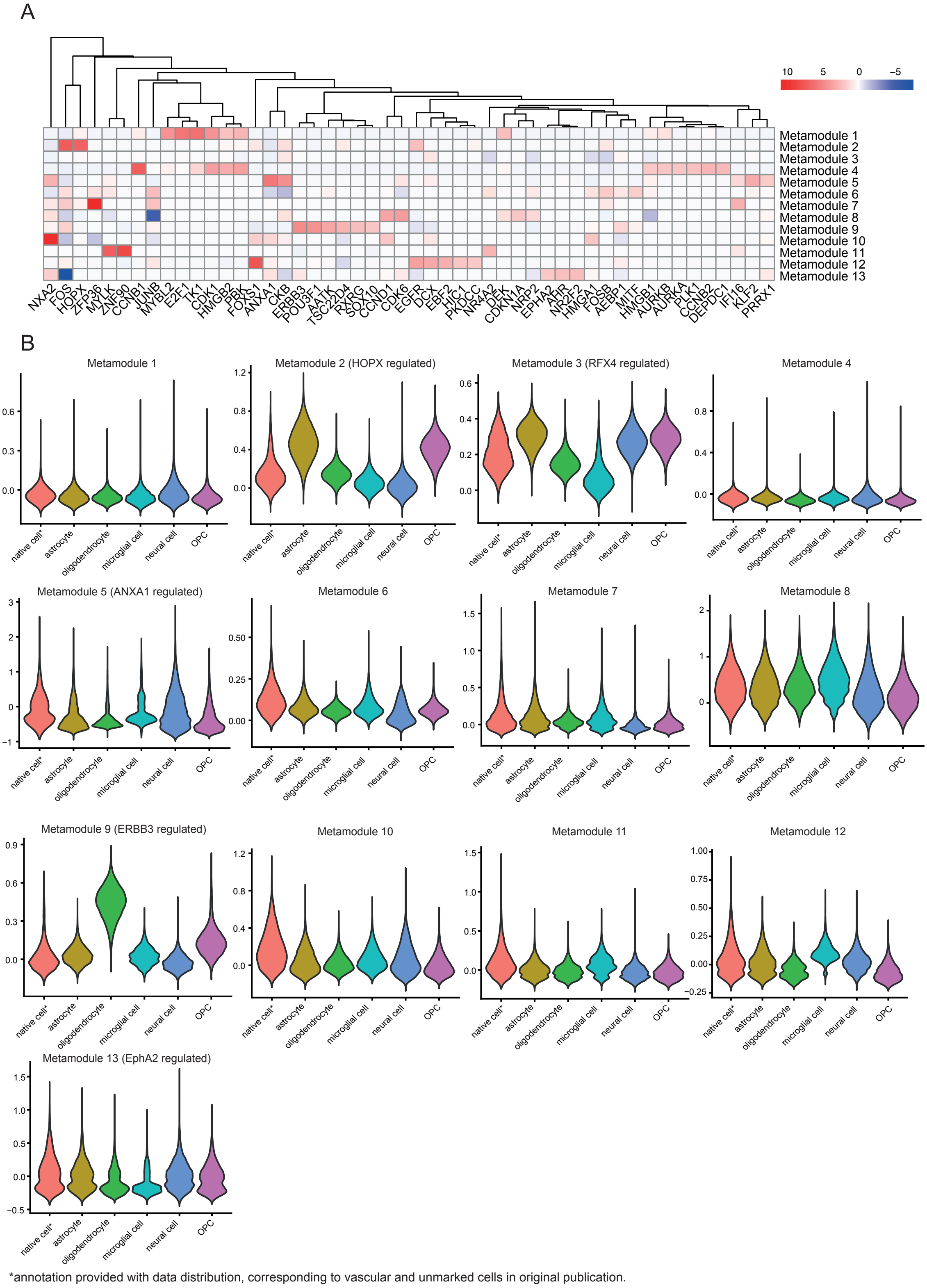


**Supplementary Figure 4: scregclust metamodules.** (A) Regulator assignments for each metamodule, relating to Figure 3 of the main manuscript. (B) enrichment of metamodule signatures in cells from [1], separated by annotated cell type.

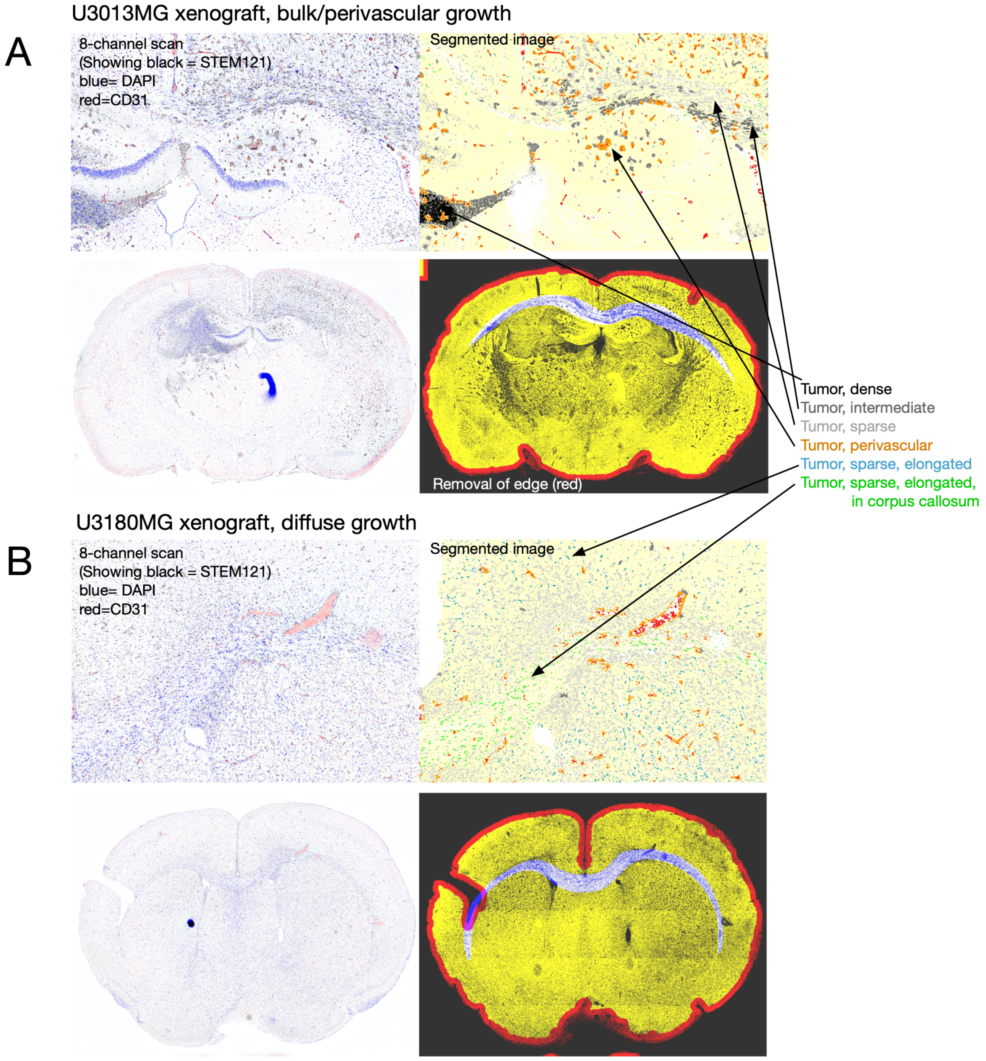


**Supplementary Figure 5: Image Segmentation:** (A) U3013MG example; (B) U3180MG example. Our image segmentation pipeline is implemented in Matlab (code available upon request). In essence, it loads a QPTIFF file representing the scan of multispectral protein staining. The pipeline begins by performing a crude segmentation using imsegkmeans (Matlab) to identify tumor, vascular, brain, and glass background pixels. Subsequently, a series of morphological operations are applied to identify each of the 9 classes described in the main manuscript. Tumor cells are delineated by watershed segmentation and assigned classes 1, 2, and 3 (as detailed in Figure 4 of the paper) based on their density. Additionally, cells are categorized as perivascular (class 4 in the main paper) if they are within 10 pixels (20 microns) of a blood vessel, as bundled tumor cells (class 5) if they are arranged in a circular pattern, detected by a Laplacian-of-Gaussian filter, and as diffuse invading cells (class 6 and 7) if they are located in low-density areas and have an eccentricity greater than 0.9. Class 6 cells are situated within the corpus callosum (manually defined for each scan), while class 7 cells are outside of the corpus callosum. Classes 8 and 9 represent blood vessels and background brain, respectively, as defined by the original imsegkmeans clustering. See also Figure 4.

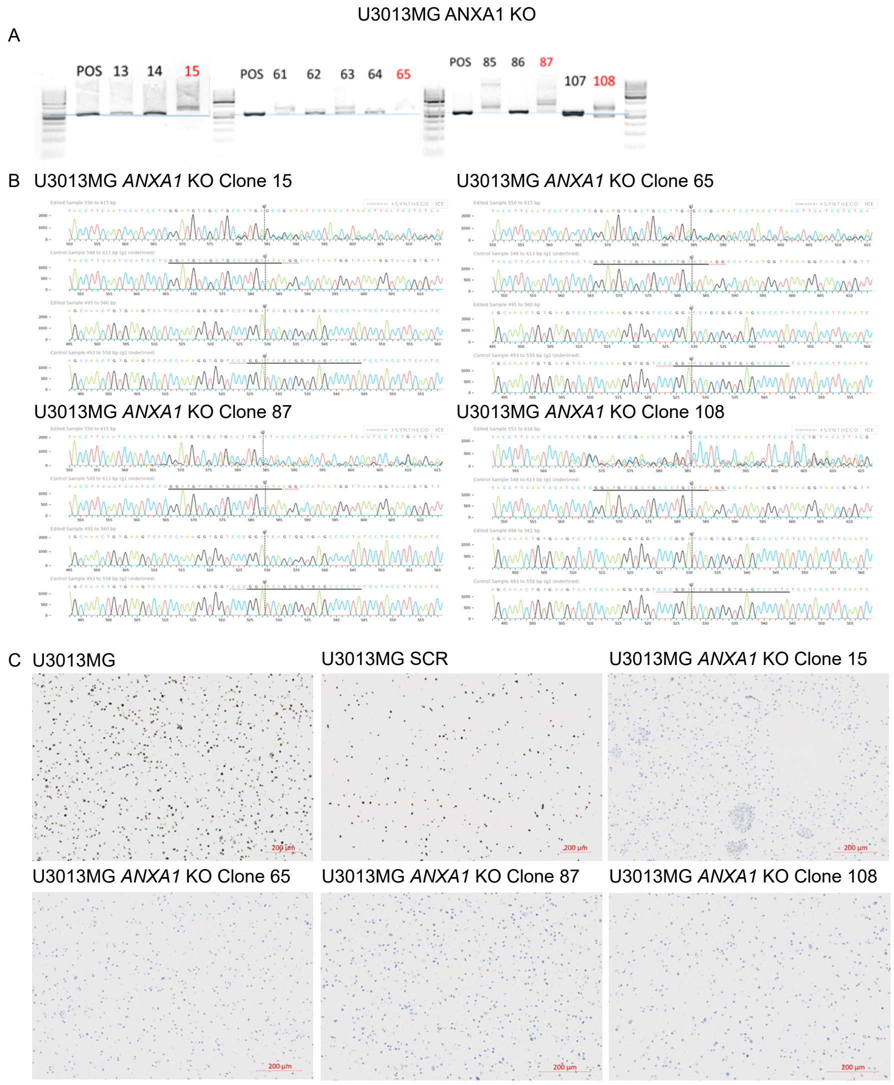


D

| Client Sample Name | U3013-luc p28 | 3013SCR | 3013 ANXA1KO-MIX1 |
|---|---|---|---|
| **Sample Code** | CL00002002 | CL00009638 | CL00009639 |
| D8S1179 | 13,14 | 13,14 | 13,14 |
| D21S11 | 31.2,32.2 | 31.2,32.2 | 31.2,32.2 |
| D7S820 | 9,11 | 9,11 | 9,11 |
| CSF1PO | 10,12 | 10,12 | 10,12 |
| D3S1358 | 17,19 | 17,19 | 17,19 |
| TH01 | 6,9,3 | 6,9,3 | 6,9,3 |
| D13S317 | 14,14 | 14,14* | 14,14 |
| D16S539 | 10,10 | 10,10 | 10,10 |
| D2S1338 | 25,26 | 25,26 | 25,26 |
| D19S433 | 14,14 | 14,14 | 14,14 |
| vWA | 16,16 | 16,16 | 16,16 |
| TPOX | 8,8 | 8,8 | 8,8 |
| D18S51 | 13,14 | 13,14 | 13,14 |
| AMEL | X,X | X,X | X,X |
| D5S818 | 12,12 | 12,12 | 12,12 |
| FGA | 18,24 | 18,24* | 18,24 |

**Supplementary Figure 6: Knockout confirmation, ANXA1** (A) agarose gel of PCR results for the targeted ANXA1 region, with alterations in clones with index 15, 65, 87, and 108. (B) Sanger sequencing confirmation of knockout in each clone. (C) ANXA1 protein staining for the source culture, scramble control (SCR) and KO clones. (D) STR profiling of the 1:1:1:1 mixture of the knockout clones which was used for the mouse experiments. See also Figure 6.

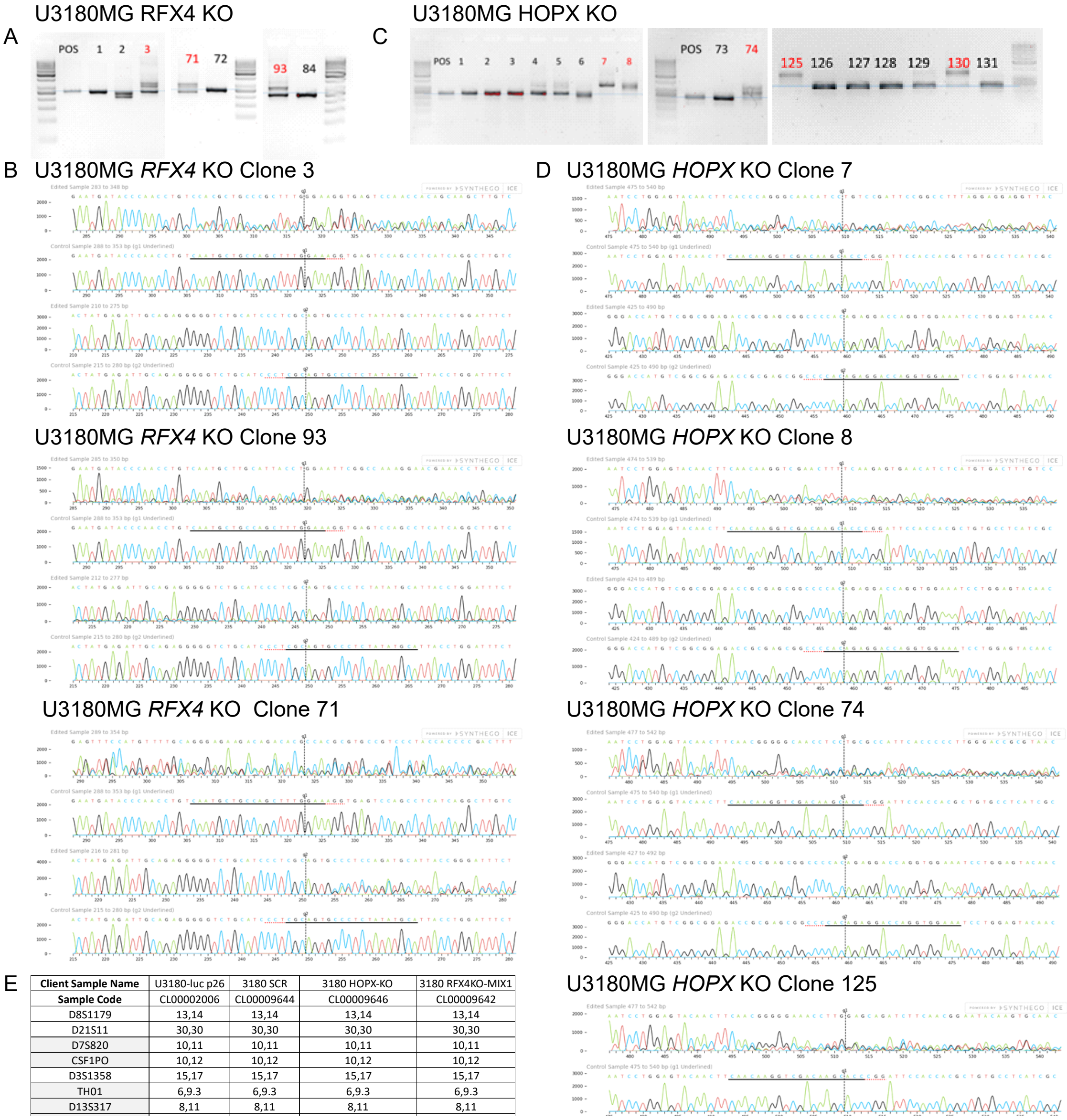


| **Client Sample Name** | U3180-luc p26 | 3180 SCR | 3180 HOPX-KO | 3180 RFX4KO-MIX1 |
|---|---|---|---|---|
| **Sample Code** | CL00002006 | CL00009644 | CL00009646 | CL00009642 |
| D8S1179 | 13,14 | 13,14 | 13,14 | 13,14 |
| D21S11 | 30,30 | 30,30 | 30,30 | 30,30 |
| D7S820 | 10,11 | 10,11 | 10,11 | 10,11 |
| CSF1PO | 10,12 | 10,12 | 10,12 | 10,12 |
| D3S1358 | 15,17 | 15,17 | 15,17 | 15,17 |
| TH01 | 6,9.3 | 6,9.3 | 6,9.3 | 6,9.3 |
| D13S317 | 8,11 | 8,11 | 8,11 | 8,11 |

**Supplementary Figure 7: Knockout confirmation** (A) agarose gel of PCR results for the targeted RFX4 region, with alterations in clones with index 3,71,93. (B) Sanger sequencing confirmation of knockout in each clone. HOPX (C) agarose gel of PCR results for the targeted HOPX region, with alterations in clones with index 7,8,74,125,130. (D) Sanger sequencing confirmation of knockout in each clone. (E) STR profiling of the uniform mixture of the knockout clones which was used for the mouse experiments. See also Figure 6.

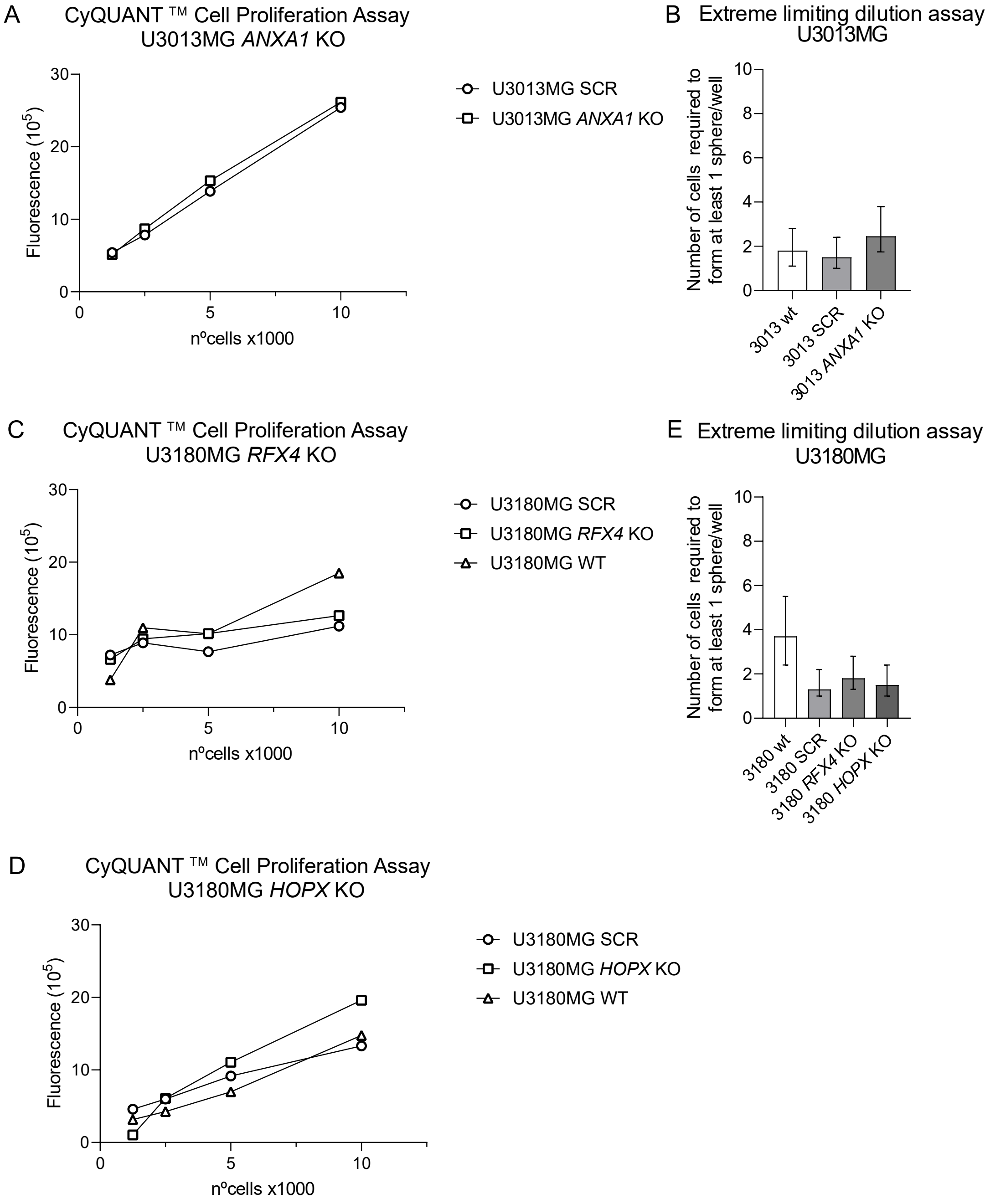


**Supplementary Figure 8: In vitro characterization of glioblastoma cells with knockout of ANXA1, RFX4, and HOPX** (A) Cell proliferation assay for U3013MG SCR and U3013MG *ANXA1* KO. (B) Extreme limiting dilution assay for U3013MG, U3013MG SCR, and U3013MG *ANXA1* KO. (C) Cell proliferation Assay for U3180MG, U3180MG SCR and U3180MG *RFX4* KO. (D) Cell proliferation Assay for U3180MG, U3180MG SCR and U3180MG *HOPX* KO. (E)) Extreme limiting dilution assay for U3180MG, U3180MG SCR, and U3180MG *RFX4* KO, and U3180MG *HOPX* KO

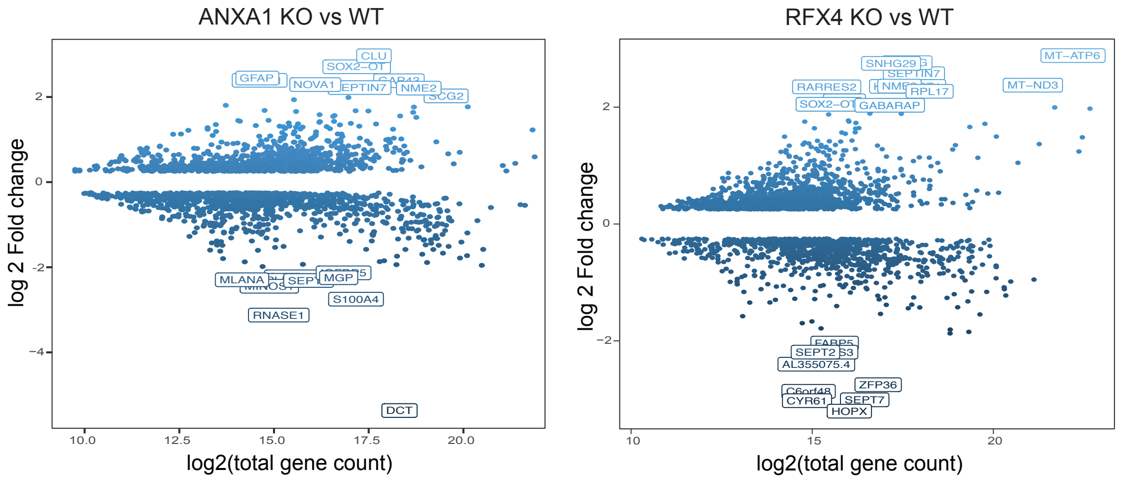


**Supplementary Figure 9: Differentially expressed genes** detected by scRNAseq of ANXA1 knock-out U3013MG PDXs compared to U3013MG (scramble gRNA) controls (left) and RFX4 knockout U3180MG PDXs compared to U3180 (scramble gRNA) controls. See also Figure 7.

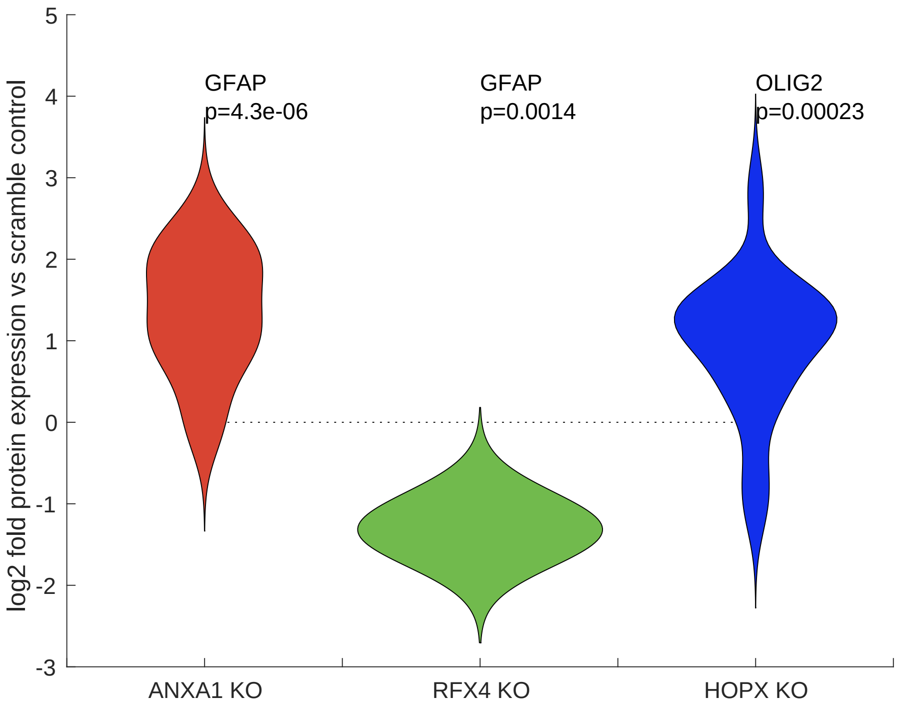


**Supplementary Figure 10: Quantitation of GFAP and OLIG2 protein staining**. A log fold change of 0 corresponds to the negative control. The red distributions illustrate the log fold values for GFAP in STEM121 positive regions in brains grafted with ANXA1 knockout U3013MG cells, compared to brains grafted with scramble control U3013MG cells. Similarly, the green distributions represent the log fold values for GFAP in STEM121 positive regions in brains grafted with RFX4 knockout U3180MG cells, compared to brains grafted with scramble control U3180MG cells. Additionally, the blue distributions depict the log fold values for OLIG2 in STEM121 positive regions in brains grafted with HOPX knockout U3180MG cells, compared to brains grafted with scramble control U3180MG cells. See also Figure 7.
